# Extracellular vesicle mobility in collagen I hydrogels is influenced by RGD-binding integrins

**DOI:** 10.1101/2024.05.29.596426

**Authors:** Nicky W. Tam, Alexander Becker, Agustín Mangiarotti, Amaia Cipitria, Rumiana Dimova

## Abstract

Extracellular vesicles (EVs) are a diverse population of membrane structures produced and released by cells into the extracellular space for the intercellular trafficking of cargo molecules. They are implicated in various biological processes, including angiogenesis, immunomodulation, and cancer cell signaling. While much research has focused on their biogenesis or their effects on recipient cells, less is understood about how EVs are capable of traversing diverse tissue environments and crossing biological barriers. Their interactions with extracellular matrix components are of particular interest, as such interactions govern diffusivity and mobility, providing a potential basis for organotropism. To start to untangle how EV-matrix interactions affect diffusivity, we use highspeed epifluorescence microscopy, single particle tracking, and confocal reflectance microscopy to analyze particle mobility and localization in extracellular matrix-mimicking hydrogels composed of collagen I. EVs are compared with synthetic liposomes and extruded plasma membrane vesicles to better understand the importance of membrane composition on these interactions. By treating EVs with trypsin to digest surface proteins, we determine that proteins are primarily responsible for EV immobilization in collagen I hydrogels. We next use a synthetic peptide competitive inhibitor to narrow down the identity of the proteins involved to argynylglycylaspartic acid (RGD) motif-binding integrins, which interact with unincorporated or denatured non-fibrillar collagen. Moreover, the effect of integrin inhibition with RGD peptides has strong implications for the use of RGD-peptide-based drugs to treat certain cancers, as integrin inhibition appears to increase EV mobility, improving their ability to infiltrate tissue-like environments.

---

Extracellular vesicles (EVs) are membrane-bound structures produced by cells and released into the extracellular space that function in cell-to-cell signaling and the trafficking of various materials.^1–5^ Biogenesis and release of EVs involves different cellular pathways, resulting in a highly heterogeneous mix of particles, from micrometer-sized microvesicles derived directly from the plasma membrane^6,7^ to 100 nm-sized and smaller exosomes produced through the endosomal pathway.^8–10^ EVs are implicated in both physiological and pathological processes, including immunomodulation,^11,12^ maintenance of pluripotency in stem cells,^13–15^ extracellular matrix (ECM) remodeling,^16,17^ and directed cell migration.^18^ Research on the functional roles of EVs has primarily been conducted *in vitro* and much focus has been placed on their role in cancer cell signaling.^2,14,19–22^ Indeed, there is growing interest in the use of EVs as prognostic markers for cancer progression.^23,24^ Despite the amount of research on their biogenesis within cells and their effects on target cell physiology, the process by which they travel between source and target remains poorly understood.^25^

Besides fundamental research in understanding their role in living systems, there is also much interest in EVs and EV-like particles as inspiration for nanomedicine and nanoparticle-based drug delivery applications.^1,26–31^ The near ubiquity of EVs in biological fluids^32,33^ implies their ability to traverse diverse tissue environments and cross different biological barriers,^30,31^ while distribution patterns of EVs injected *in vivo*^34^ suggest some degree of tissue specificity – two characteristics that would be highly advantageous for the targeted delivery of therapeutics. The way EVs interact with different ECM materials is thus an important research topic, as these interactions likely form the basis for organotropism and tissue specificity.^35,36^

Previous studies have determined that enzymes and receptors on EV surfaces are active and capable of binding to or interacting with ECM components using biochemical pulldown assays or analyses of substrate species on the molecular level.^16,37–39^ Certain studies have also characterized *in vivo* systemic EV behaviour and organotropic distribution arising from expressed surface proteins.^16,40,41^ However, a micro-to mesoscale link between individual molecular interactions and their functional consequences on EV behaviour as a whole is missing. To address this, we aim to link the molecular interactions at the EV surface with the ECM and their consequences in EV diffusion and mobility. Our hypothesis is that EV mobility and their ability to infiltrate tissues are governed by surface interactions, and that this will be reflected in their Brownian motion as they interact with their molecular environment. Differences in mobility at this level may eventually scale up to differences in tissue infiltration as they interact with the myriad molecules that make up the tissue microenvironment, leading to systemic specificity and homing behaviour.

Here, we investigate EV interactions in hydrogels composed of collagen I. We first collect and purify EVs from a breast cancer cell line using size exclusion chromatography (SEC)^42,43^ and characterize them using LAURDAN fluorescence spectroscopy and Western blot analysis of proteins. EV mobility in collagen gels is studied using single particle tracking^44^ on image sequences obtained with highspeed epifluorescence imaging of EVs introduced into preformed collagen gels. EV behaviour is compared to that of synthetic liposomes, as well as vesicles composed of extruded whole-cell plasma membrane to better understand the influence of EV membrane composition on mobility. We also analyze particle localization in collagen gels relative to fibrils imaged with confocal reflectance microscopy. Finally, we determine that integrins are primarily responsible for dictating the mobility of EVs by treating EVs with trypsin to digest surface proteins and with a competitive inhibitor peptide to specifically block integrin-collagen interactions. With our results, we show that EV membrane composition imparts specificity in ECM interactions that affect not only overall mobility, but also spatial distribution within a collagen I matrix.

## Results and Discussion

### EVs and plasma membrane vesicles: collection, synthesis and characterization of size and composition

EVs were collected from cultured MDA-MB-231 breast cancer cells, which have previously been used in EV research.^45^ Cells were incubated over three days with serum-free media to avoid contamination with exogenous vesicle material. EVs released into the conditioned media were collected and purified using SEC.^42,43,46^ Approximately 20 fractions of 500 μL volume were separated and collected. Analysis of SEC fractions with dynamic light scattering (DLS) showed the highest enrichment of 100-400 nm diameter particles, assumed to be EVs, in fractions 7-10 (see Supplemental Fig. S1).

It is well-documented that EVs have a distinct membrane composition compared to the plasma membrane of their source cell, and that this also varies between EV subpopulations,^7,47^ both in terms of enriched lipid species and proteins. To better understand the functional effects of such differences, we produced vesicles that should more closely reflect the composition of whole cell plasma membrane while being approximately the same size as the collected EVs (Fig. 1; see Suppl. Fig. S1 for size distributions). Giant plasma membrane vesicles (GPMVs) were first produced according to a previously published protocol by exposing cells to N-ethylmaleimide (NEM) as a vesiculation agent.^48^ This protocol resulted in blebs with cell-like mechanical properties^49^ that were then extruded to produce 200 nm-diameter ‘large’ plasma membrane vesicles (LPMVs).^50^

**Figure 1.**
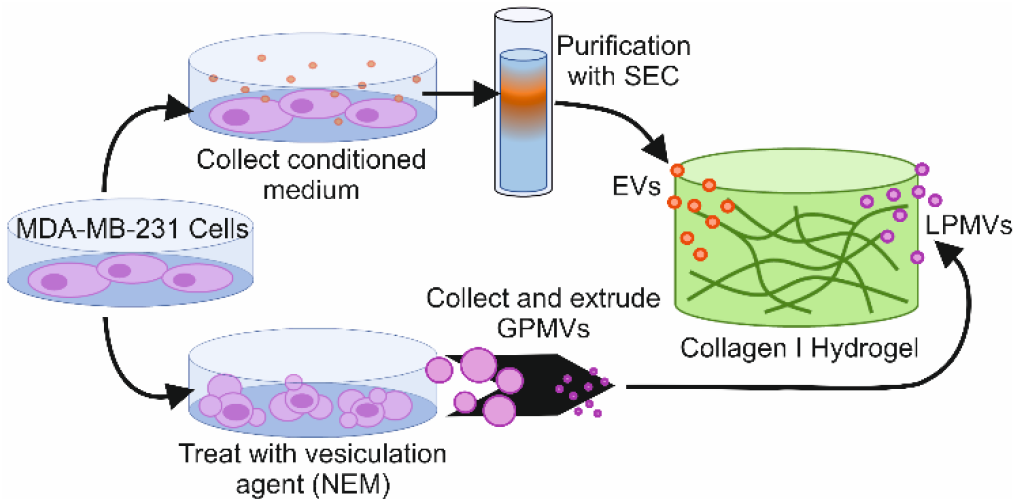
Schematic diagram of experimental approach to produce EVs and LPMVs. EVs are collected from the supernatant of cultured MDAMB-231 cells and are purified with SEC (upper path). LPMVs are generated by treating cells with NEM as a vesiculation agent, collecting the formed GPMVs, and extruding them into EV-sized LPMVs (lower path). Particles are allowed to diffuse into fully-formed collagen I hydrogels and are imaged with confocal microscopy or epifluorescence microscopy.

Figure 2A shows cryogenic scanning electron microscopy (cryoSEM) images of EVs and LPMVs appearing as round structures with sizes corresponding to the size distributions measured with DLS (See Suppl. Fig. S2, S3 for further images). Samples were high-pressure frozen in disks, which were cleaved laterally in half to expose the particles embedded in the surface of the surrounding frozen medium. EVs appear to have a size range between 100 nm and 400 nm in diameter, while LPMVs appear approximately 200nm in diameter and below. The existence of smaller particles <100 nm in our LPMV samples makes sense, as there was no lower limit on the size of the LPMVs during extrusion, nor were the particles purified as the EVs were with SEC. They likely did not show up in our DLS measurements due to being below the limit of detection for the device used.

**Figure 2.**
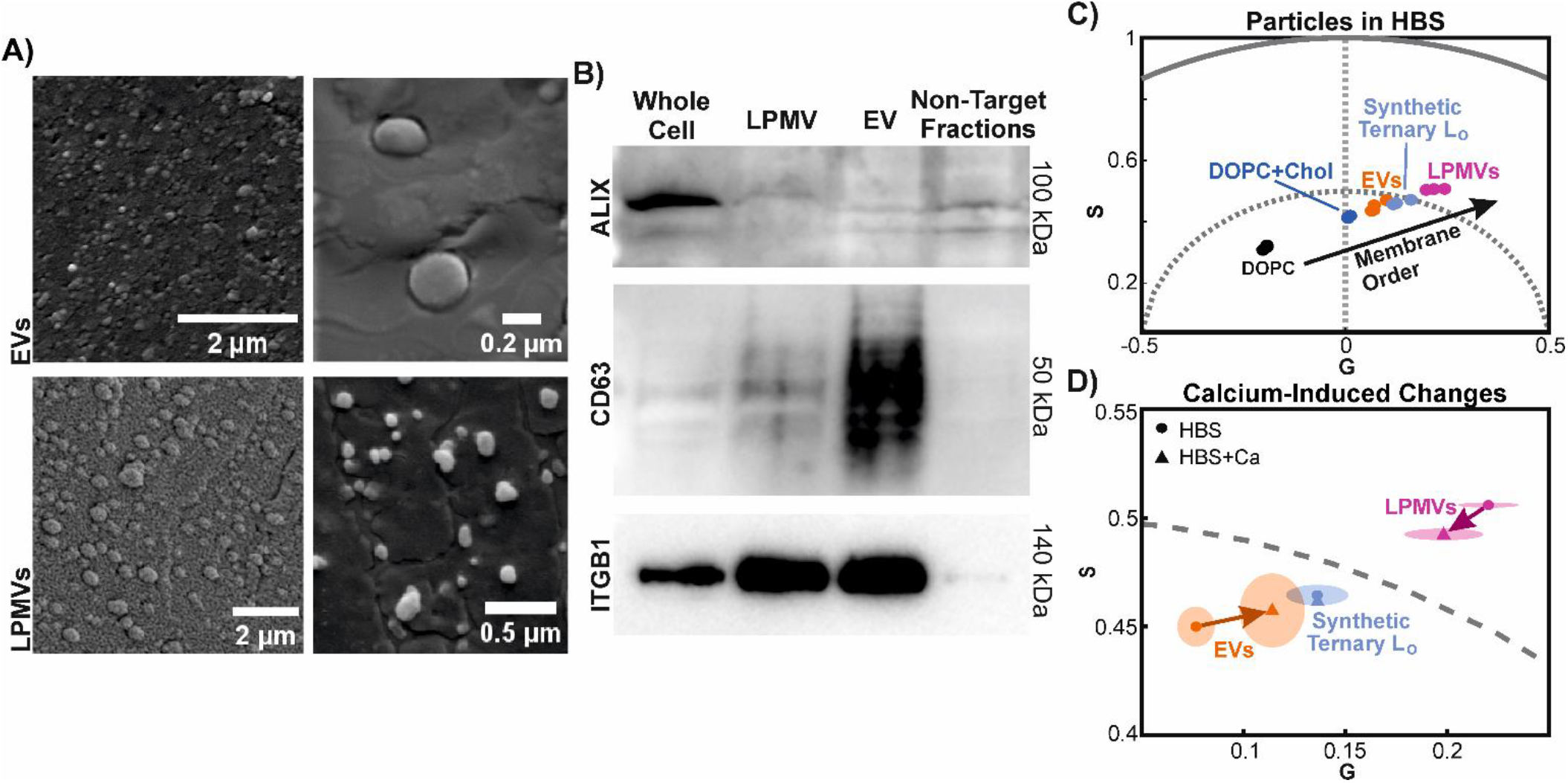
Characterization and comparison of EVs and LPMVs. A) Representative cryoSEM images of EVs and LPMVs. See Suppl. Fig. S2, S3 for more examples. Their apparent sizes correspond to size distributions obtained with DLS (see Suppl. Fig. S1). B) Western blot array characterizing, from left to right, relative protein expression in whole cell lysates (positive control), LPMVs, EVs, and the non-EV fractions collected during SEC purification of EVs (negative control). Whole protein concentrations of samples were determined with a Bradford assay and the samples were diluted such that each lane in the blot contains the same whole protein content. Apparent sizes of the probed proteins according to their position in the gel are shown on the right. Densitometric analysis and the uncropped blots are shown in Suppl. Fig. S4. C) Spectral phasor analysis of LAURDAN fluorescence spectroscopy data of EVs, LPMVs, and synthetic multilamellar vesicles (DOPC, DOPC+Chol, Synthetic Ternary Lo) in calcium-free HEPES buffered saline (HBS). The synthetic lipid vesicles form a linear trajectory on which molecular packing, membrane dehydration, and membrane order are expected to increase along the linear trajectory indicated by the black arrow. Since EVs and LPMVs fall on this alignment, the state of molecular packing and hydration in their membrane bilayers can be approximated with minimal synthetic lipid mixtures, with EVs having an intermediate state between DOPC+Chol and the synthetic ternary Lo mixture. LPMVs lie beyond the ternary Lo mixture, suggesting a greater degree of packing and membrane order. Individual points represent separate experimental replicates. D) Shifts in phasor position due to the presence of calcium. Markers indicate the centre of mass of data points for membranes in calcium-free (HBS; circles) and calcium-containing (HBS+Ca; triangles) buffers. Shaded areas represent the standard deviation of the Cartesian coordinates, G and S of the spectral phasor plot (see Eqs. 1 and 2 for definition) from n=3 experimental replicates. Arrows indicate the direction of the shifts in centre of mass of the data points. Calcium causes increased packing in EVs and decreased packing in LPMVs, emphasizing the differences in their membrane compositions and possibly also their interactions with the environment. Shifts of such magnitude are not observed in the synthetic lipid vesicles.

To confirm that our collected EVs express standard EV markers and to better differentiate whether they are exosomes or microvesicles, we used Western blot to probe for commonly expressed markers (Fig. 2B; Suppl. Fig. S4). Particle size is often correlated with EV subgroup and mechanism of biogenesis and release, with microvesicles being, on average, larger than EVs.^5,7^ Natural size variation also occurs within subgroups, however, and the size range of our collected EVs reflects the natural size variation in cell-derived vesicles. Western blot thus remains the most effective way to differentiate between EV subtypes. We compared expression of these markers to that of our LPMVs, as well as whole cell lysates as a positive control, and the non-EV fractions obtained with SEC as a negative control. These non-target SEC fractions contain particles smaller and larger than the target range of 100-400 nm and most likely consist of cellular debris and free proteins in suspension, as well as EVs or EV aggregates. Samples were diluted and normalized to have the same whole protein content in each lane. We first investigated ALG-2-interacting protein X (ALIX), an accessory protein of the endosomal sorting complexes required for transport (ESCRT) that is involved in the packaging of specific cargo into EVs, as well as their formation *via* multivesicular bodies (MVBs) and the endosomal system^8,51–53^. We also investigated CD63, a tetraspanin commonly used as a specific marker for exosomes originating from endosomal structures^54,55^. Finally, we probed the expression of integrin-β1 (ITGB1) as a surface protein commonly found in EVs^36^ that could also have a functional role in binding to ECM components. ALIX is expressed at a high level in our whole cell lysates, appearing as a band at 100 kDa and suggesting active MVB formation. Expression appears relatively low elsewhere, with double bands visible in the negative control and EV samples and a possible single band appearing in the LPMV sample. This would suggest that ALIX expression continues on in nontarget SEC fractions, likely representing smaller EVs than our 100-400nm diameter target fractions. Similar differential expression of ALIX has previously been reported in EVs from the MDA-MB-231 cell line by Kong et al.^56^ Furthermore, they also reported relatively low ALIX expression in EVs compared to whole cell lysates, at least in their control condition, similar to our result. The low expression of ALIX in our EVs is likely due to the much higher relative expression of other proteins, such as CD63 and ITGB1. ALIX is evidently still expressed, but appears less enriched than other markers after normalizing to whole protein content. A similar effect can be found in the Western blot data of González-King et al.,^45^ where relatively low ALIX expression can be seen in EVs from MDA-MB-231 cells alongside high expression of flotillin and CD63. Using densitometry to analyze our Western blot data (Suppl. Fig. S4), ALIX expression appears to be inversely proportional to CD63 and ITGB1 expression across different samples, further supporting this possibility. CD63 is highly enriched in the target EV fractions and virtually nonexistent in the off-target fractions, appearing as a smear of bands between 25-60 kDa. This smearing is consistent with previous reports and is due to variable glycosylation of the protein, which can affect migration during electrophoresis.^57^ Meanwhile, low ALIX and CD63 expression in LPMVs reflects their origin as artificial plasma membrane structures. Finally, ITGB1 is enriched in both EVs and LPMVs compared to whole cell lysates, and is not expressed in the off-target negative control. Altogether, this would suggest that our SEC-purified EVs largely represent CD63-positive, MVB-originating exosomes. Our LPMVs, meanwhile, would be more representative of outer plasma membrane, lacking markers of endosomal origin but having high ITGB1 expression. While microvesicles derive from direct pinching off of the plasma membrane, our LPMVs likely are not fully representative of them, since microvesicles are specialized structures that arise through specific membrane processes.^5,6^ LPMVs would thus lack the enrichment of molecules that are involved in their biogenesis and release and would more closely resemble whole plasma membrane.

To further our understanding of the biophysical consequences of different membrane compositions, we conducted LAURDAN fluorescence spectroscopy to probe membrane phase state and lipid packing in EVs and LPMVs (Fig. 2 C,D). LAURDAN is a lipophilic fluorescent probe whose emission spectrum is sensitive to the molecular environment of the lipid bilayer into which it is inserted.^48,58,59^ Differences in membrane phase state, hydration, or lipid packing can be represented in graphical form using phasor analysis, which decomposes the spectral shifts into the Cartesian coordinates, G and S.^60–64^ For comparison and to better understand the differences between EVs and synthetic systems, we have also analyzed synthetic multilamellar lipid-only vesicles composed of pure 1,2-Dioleoyl-sn-glycero-3-phosphocholine (DOPC); a binary mixture of 70% DOPC and 30% cholesterol (DOPC+Chol); and a ternary mixture of 13% DOPC, 44% Dipalmitoylphosphatidylcholine (DPPC), and 43% cholesterol (Ternary L_o_). While DOPC and DOPC+Chol represent lipid membranes in the liquid-disordered (L_d_) phase state, the Ternary L_o_ mixture represents a liquid-ordered lipid membrane, referring to the molecular packing in the lipid bilayer and order in the hydrophobic fatty acid tails.^65^ These synthetic lipid vesicles fall in a linear trajectory on the phasor plot, which represents increasing membrane packing from one end to the other. EVs fall between DOPC+Chol and the Ternary L_o_ mixture, suggesting an intermediate liquid-ordered phase state. LPMVs, however, fall beyond the Ternary L_o_ mixture on the same alignment, suggesting a greater degree of membrane packing and order. The fact that both EVs and LPMVs are aligned with the axis of the synthetic lipid mixtures suggests that the state of molecular packing and hydration in their bilayers can be mimicked with minimal lipid membrane structures. In this regard, it has previously been reported that other biophysical properties of cell-derived vesicles, such as viscosity and mechanical stiffness can be reconstituted with synthetic liposomes.^66^

Both types of cell-derived membranes appear to be responsive to the presence of calcium. Adding 2mM calcium to the medium results in an increase in packing in EVs and a decrease in LPMVs (Fig. 2D). Shifts of such magnitude in phasor position do not occur in the synthetic DOPC, DOPC+Chol, and Ternary L_o_ mixtures. These calcium-induced changes may be related to the presence of other, possibly charged lipid species or to the presence of proteins or glycocalyx. Our vesicles lack charged phospholipids, such as phosphatidylserine, which is enriched in EVs^47^ and cause calcium ions to localize closer to the membrane.^67^ We note, however, that the overall zeta-potential of EVs and LPMVs is similar to that of pure DOPC large unilamellar vesicles (LUVs; see Suppl. Fig. S5). The opposing effects of calcium on EVs and LPMVs also emphasizes the compositional and potentially functional differences between them.

### EV mobility in collagen hydrogels

With the differences in membrane composition observed between EVs and LPMVs, we wanted to investigate if and how these differences would be reflected in their interactions with ECM materials, and how this might functionally affect their diffusion through an ECM-like environment. In light of the highly complex molecular environment of tissues, we used collagen I hydrogels to better systematically and quantitatively study particle diffusion in a model hydrogel environment. Collagen I is one of the most abundant proteins in mammalian ECM and plays an important structural role in tissues, as well as providing contextual signaling cues to cells.^68–70^ Moreover, collagen is often used in tissue engineering applications as cell scaffolds.^64,71,72^

To study the mobility of EVs and LPMVs in collagen I hydrogels, gels were first produced in the wells of a 96-well plate at a concentration of 1.5 mg·mL^-1^ in calcium-free HEPESbuffered saline (HBS) and calcium-containing HEPES buffered saline (HBS+Ca). Gels were allowed to fully polymerize at 37°C overnight. Figure 3A shows a representative fully-formed fibrillar collagen I matrix imaged with confocal reflectance microscopy. To characterize the hydrogels, the mesh size was estimated using a previously described protocol^72,73^ with slight modification. Confocal reflectance images were first binarized and skeletonized (Fig. 3B) to remove scanner artifacts and the sizes and number of spaces between fibrils in the x- and y-directions were counted up. These fell into an exponential distribution (Fig. 3C), whose mean value was determined by fitting an exponential function to the data. This mean mesh size does not appear to be sensitive to the presence of calcium in the buffer (Fig. 3D).

**Figure 3.**
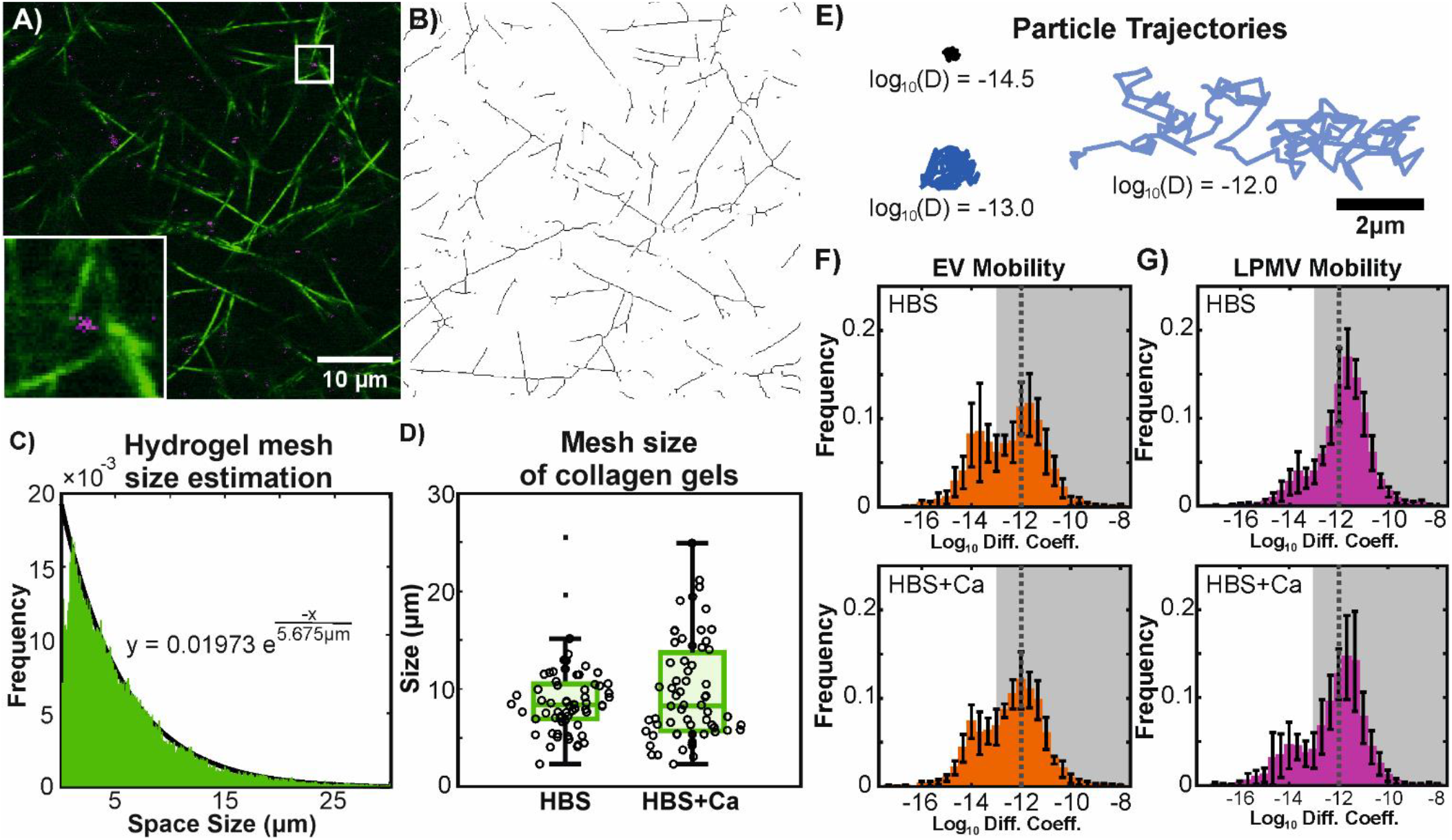
EV and LPMV diffusion in collagen I hydrogels. A) confocal reflectance micrograph showing collagen fibrils in green and fluorescentlylabelled EVs in magenta. Bottom left inset shows a particle in the indicated region zoomed in and corresponds to a 5×5µm region. B) The image in (A) is binarized and skeletonized in order to determine mesh size. C) Mesh size is estimated by counting up the number and sizes of spaces in the x- and y-directions of binarized images. In histogram form, the sizes of the spaces fall into an exponential distribution, where the mean value is the reciprocal of the exponential coefficient of a fitted exponential function. D) The mesh size of collagen gels is not significantly affected by the presence of calcium, as determined with a 2-way T-test (p>0.05), but varies greatly between and within samples. Circles represent individual samples. Dots represent outlier data that were excluded from the rest of the box plot. E) Example particle trajectories showing an immobile particle (green, log10(D)=-14.5), a particle at the mobility cutoff (blue, log10(D)=-13.0) and a highly mobile particle with the Stokes-Einstein-predicted diffusion coefficient for a 200nm particle diffusing in liquid water (magenta, log10(D)=-12.0). F,G) histograms of log10 diffusion coefficients of EVs (F) and LPMVs (G) diffusing in collagen hydrogels formed without (upper row; HBS) and with (lower row, HBS+Ca) calcium present in the buffer. Error bars show standard deviation over n=7 (EVs) or 8 (LPMVs) experimental replicates. The mobile fraction is represented by the shaded grey area with the cutoff at -13.0. A dotted line shows the Stokes-Einstein prediction for an ideal 200nm particle diffusing in liquid water. Mobile fraction and overall shape of distributions for both particles appear insensitive to calcium, but LPMVs are clearly more mobile than EVs.

Particles were introduced to the gels by pipetting suspensions directly on top of gels and allowing them to diffuse throughout for at least three hours prior to imaging. A highspeed camera mounted to a standard light microscope in epifluorescence mode was used to image the particles as they were diffusing. Single particle tracking was conducted on acquired image sequences using the MOSAIC suite plugin developed for FIJI by Sbalzarini et al.^44^ Representative particle trajectories are shown in Figure 3E, exhibiting a wide range of possible particle mobilities. To better visualize the differences in particle mobilities, the base-10 logarithms of the diffusion coefficients obtained from particle tracking are shown. Histograms of these values (Fig. 3F,G) show strong bimodal behavior, similar to what has been previously reported for synthetic LUVs and polymeric nanoparticles in various hydrogel environments.^74–76^ The peak appearing near the value of -12 represents mobile particles approaching the Stokes-Einstein prediction for the diffusion coefficient of an ideal 200 nm-diameter spherical particle diffusing in liquid water at room temperature. The other peak at -14 would then represent immobilized particles, with its mean value linked to the image resolution and frame rate of our acquisition setup, as well as the apparent particle size as they appear in our images.^74^ To quantify particle mobility, we defined the value of -13.0 as a cutoff point that separates well the two peaks in the distributions, and summed the bins of the histograms that fall above this cutoff to obtain a mobile fraction. In previous work involving synthetic LUVs diffusing in agarose hydrogels,^74^ we were able to define a mobile cutoff based on the lower limit of detection of particle movement of our imaging setup. This was not possible here due to the polydispersity of our EVs making the apparent size of particles in images poorly-defined. Despite this, with the current mobility cutoff of -13.0, it is clear that LPMVs are significantly more mobile than EVs, and that the mobilities of both particles appear unaffected by the presence of calcium.

The variation in particle size of EVs and LPMVs is unlikely to play a major role in mobility. Since the average mesh size of the collagen gels is on the order of several micrometers and above while the particles are, at most, 400 nm in diameter, steric effects are unlikely to affect particle diffusion. We have previously reported on this topic using synthetic liposomes diffusing in agarose hydrogels.^74^ Furthermore, although the Stokes-Einstein prediction for an ideal spherical particle’s diffusion coefficient is inversely proportional to the particle’s size, we note that the difference between the base-10 logarithms of diffusion coefficients of particles with 100 nm and 400 nm diameters (our maximum size range) comes out to be 0.6, which is far smaller than the full range of values we measure, from -18.0 to -8.0. Thus, the differences in particle mobility we observe must arise from other causes.

It is likely that the observed differences in particle mobility is due to differences in membrane composition between EVs and LPMVs. It must be noted, however, that the vesiculation agent, NEM can irreversibly react with and modify cysteines and thiol groups,^48,77^ leading to changes in the membrane constituents of LPMVs. Despite this drawback, the use of NEM to produce plasma membrane vesicles seems a better option for maintaining protein integrity compared to other protocols that use paraformaldehyde and dithiothreitol as vesiculation agents,^50,78^ which would result in protein crosslinking and fixation. While protocols that do not use such harsh chemicals exist, based on the use of salt buffers to induce osmotic shocks,^79^ such protocols take significantly more time and produce less membrane material, which is critical to our studies due to sample loss during extrusion.

### Mimicking EV diffusion with synthetic vesicles

To further our understanding of the interactions between EVs and collagen I, we used synthetic LUVs to see if our observed particle behavior could be reproduced using minimal model membranes. As a first attempt, we produced 200 nm-diameter DOPC LUVs (PC-LUVs) and added them to fully-formed collagen hydrogels with calcium-free (HBS) and calcium-containing buffer (HBS+Ca), in both of which they were observed to become fully immobilized (Fig. 4A; see Suppl. Fig. S6 for histograms of log_10_ diffusion coefficients). This adhesion between DOPC and collagen I appears to be particularly strong, as our previous work has shown that DOPC LUVs remain relatively mobile in agarose hydrogels despite steric trapping effects from the much smaller pore sizes.^74^ Our collagen I hydrogels, meanwhile, have a mesh size on the order of several micrometers (Fig. 3D) – sufficiently large relative to the particle diameter that diffusing particles should be unaffected by steric trapping effects. We next explored whether including 1,2-dioleoyl-sn-glycero-3-phospho-L-serine (DOPS) as a negatively charged phospholipid upregulated in EVs would improve mobility. Phosphatidylserines have previously been found to comprise up to 20% of EV membranes from various cell sources,^7,47,80^ so we produced PS-LUVs comprised of a 4:1 molar ratio mixture of DOPC and DOPS. PS-LUVs are fairly mobile in calcium-free buffer, but become immobilized in calcium-containing buffer, likely due to electrostatic interactions and calcium ‘bridging’^81,82^ between the phosphatidylserine groups and the matrix collagen. We then explored how PC-LUVs could be modified to rescue their mobility. We tried ‘blocking’ their surfaces by incubating them with soluble collagen I (bPC-LUVs) or by including PEGylated lipids (PEG-LUVs) into their membranes. Blocking with 0.05 mg·mL^-1^ collagen I significantly increases the mobility of PC-LUVs, suggesting that simply having a buffer layer to prevent direct adhesion of the matrix collagen to the phospholipid membrane surface is enough to improve particle mobility. Inclusion of 1 mol% 1,2-distearoyl-sn-glycero-3-phosphoethanolamine-N[methoxy(polyethylene glycol)-5000] (DSPE-mPEG5K) is more effective at preventing adhesion and restores the mobile fraction of PC-LUVs to a value similar to that of EVs. The reason why mobility is not restored to an even higher fraction in the PEG-LUVs is likely due to low polymer coverage and the PEG chain being in the mushroom conformation at this concentration in the membrane,^83–85^ with parts of the underlying DOPC membrane being exposed. While PEG is often thought to improve the diffusive properties of nanoparticles by sterically preventing the adsorption of colloidal proteins,^75,84,85^ it appears that this may not entirely be the case here. It is possible that colloidal collagen molecules that have not been incorporated into fibrils exist in our hydrogels within the mesh spaces, but as was seen in the bPC-LUVs, having a layer of adsorbed collagen appears to increase mobility by preventing adhesion to some degree. It seems more likely that the PEG layer is directly preventing adhesion to the collagen fibrils.

**Figure 4.**
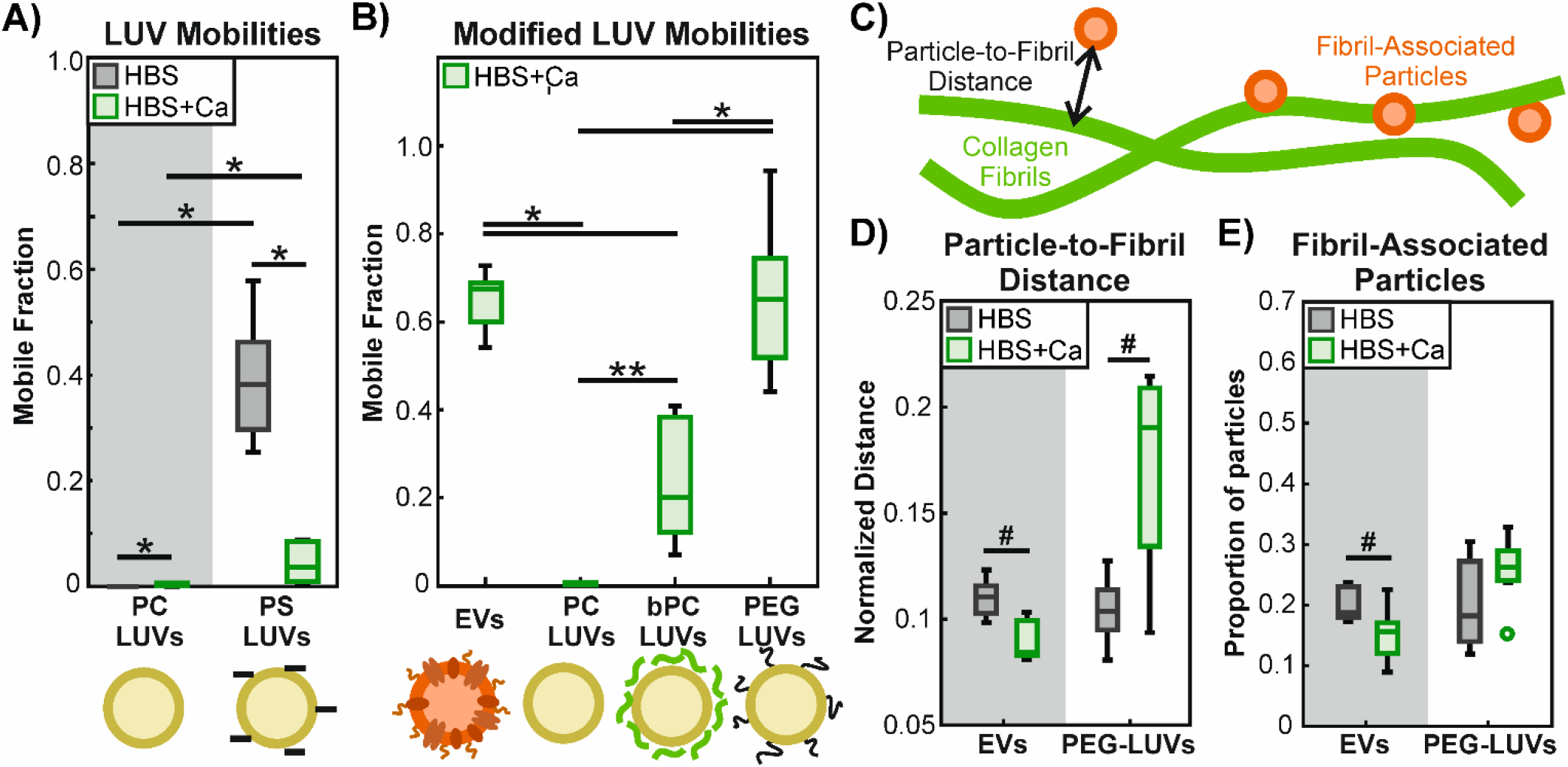
Modeling EV mobility in collagen I hydrogels with synthetic LUVs. Single particle tracking is used to determine the diffusion coefficients of imaged particles. The base-10 logarithms of the diffusion coefficients are binned into a histogram and the mobile fraction is determined by summing up the bins that lie above the mobility threshold of -13. A) The mobility of LUVs composed of pure DOPC (PC-LUVs) is compared to that of LUVs composed of 4:1 DOPC/DOPS (PS-LUVs) in calcium-containing (HBS+Ca) and calcium-free (HBS) buffers. PC-LUVs remain immobile in collagen I gels in both buffers. PS-LUVs appear fairly mobile in HBS, but become immobilized in HBS+Ca, likely due to electrostatic interactions. Significant differences are determined by 2-way ANOVA with Tukey-Kramer post-hoc analysis and are indicated by * (p<0.01) with n=6 replicates. B) EVs are compared with synthetic PC-LUVs previously incubated with 1.5 mg/mL soluble collagen I to block their surfaces (bPC-LUVs), as well as PEGylated LUVs composed of DOPC + 1 mol% DSPE-mPEG5K (PEG-LUVs) in calcium-containing buffer (HBS+Ca). Cartoons underneath illustrate the differences between the particles, with an EV depicted in orange with complex composition, synthetic LUVs depicted in yellow with negative charges from DOPS represented as minus symbols, colloidal adsorbed collagen I depicted in green, and anchored PEG chains in black. Significant differences are determined with 1-way ANOVA with Tukey-Kramer post-hoc analysis, as indicated with * (p<0.01) or ** (p<0.05). Statistics were determined with n=6 replicates. C) Schematic depiction of particle-to-fibril distance and fibril-associated particles. Particles localized within 500nm of the central axis of a collagen fibril, roughly the noise floor for determining colocalization in our collected images, are considered fibril-associated by proximity. D) Average distance from PEG-LUVs and EVs to the nearest collagen fibril, normalized by hydrogel mesh size, as measured in calcium-free (HBS) and calcium-containing (HBS+Ca) buffer. E) Proportion of PEG-LUVs and EVs determined to be associated with collagen fibrils by proximity. In D and E, no significant differences were found with 2-way ANOVA (p>0.05), but comparisons with 2-way T-tests on isolated data were found to be statistically significant for EVs, as indicated by # (p<0.01). Differences for PEG-LUVs were not significant (p>0.05) Statistics were determined with n=6 replicates.

We further investigated the interactions between PEG-LUVs and collagen I by analyzing the same samples used for single particle tracking with confocal microscopy. Collagen I fibrils were imaged in reflection mode^72,73^ and the labelled LUVs were imaged with standard fluorescence confocal microscopy. Images of fibrils and particles were binarized and the fibrils skeletonized (Fig. 3A,B) to remove pixel spread due to their widths being below the optical diffraction limit. The coordinates of the geometric centres of the particles were determined and the distance to the nearest fibril was measured using the Python implementation of the KD-Tree nearest-neighbour algorithm.^86^ We found that the average particle-to-fibril distance for each replicate was dependent on the overall hydrogel mesh size of the replicate, and that the mesh size varied greatly between samples due to natural sample heterogeneity (Suppl. Fig. S7). Alongside measurements of particle-to-fibril distance, we also determined the mesh size of each sample. By normalizing the average particle-to-fibril distance in each sample to its corresponding mesh size, we were able to obtain particle positions relative to the mesh structure (Fig. 4B), which greatly reduced the variance of the data. We found that the presence of calcium in the buffer (HBS+Ca) resulted in a greater normalized particle-to-fibril distance than particles in calcium-free buffer (HBS) for PEG-LUVs (Fig. 4C). The opposite is true for EVs.

We next defined a fibril-associated particle as being one whose centre is localized within 500 nm of the central axis of a collagen fibril (after skeletonization; Fig. 4D). This corresponds roughly to a distance of 5 pixels in our images, and thus, to the noise floor for determining colocalization: half the apparent width of the collagen fibrils plus the apparent radius of the imaged particles. For PEG-LUVs, the proportion of fibril-associated particles appears insensitive to calcium despite the increase in normalized particle-to-fibril distance. This would suggest that the affinity of the PEG-LUVs for fibrillar collagen I remains unchanged, though they tend to localize more towards the interior of the hydrogel mesh spaces when they are not sticking to the fibrils. In contrast, particle-to-fibril distance and fibril-association in EVs both decrease in the presence of calcium. This seems counter-intuitive, but may be due to an increase in transient or weakly-binding interactions, leading to EVs being closer to fibrils overall, but not necessarily colocalizing with them. There is also the possibility of non-fibrillar colloidal collagen existing within the hydrogel mesh spaces, which the EVs may be interacting with.

Because the mobile fraction and proportion of fibril-associated particles are independent measurements from separate analyses, it is not possible to comment on whether these populations overlap. Nevertheless, we can think of these values as fractions of the whole population of particles in a sample. The mobile fraction for EVs has a median value of 0.6 and the proportion of fibril-associated EVs hovers around 0.2, leaving a minimum of a fifth of particles overall, or half of the immobile fraction unaccounted for that are immobilized, but not bound to fibrils. These EVs could be binding to and becoming immobilized by unincorporated or denatured collagen molecules that cannot be imaged with confocal reflectance microscopy. Calcium could thus be altering the relative affinities that particles have for fibrillar versus colloidal or denatured collagen.

Altogether, it is evident that the behaviour of EVs can only be partially reproduced with synthetic LUVs. The complex membrane composition of EVs prevents complete adhesion to collagen while also enabling immobilizing interactions. While LPMVs also possess a complex membrane composition, differences in composition appear to result in functional differences in their mobility compared to that of EVs. This suggest a degree of specificity in these interactions that warrant further investigation.

### Integrins are responsible for EV-matrix interactions

To determine which membrane component, if any, is primarily responsible for the interaction of EVs with collagen I, we treated EVs with trypsin (tEVs) to digest their surface proteins. In doing this, we could determine if the immobilization of EVs is due to a protein while lipid species would remain intact. The glycocalyx would also be largely intact, though the digestion of glycosylated proteins may release sugar chains anchored in this way. We found a significant increase in the mobile fraction and average particle-to-fibril distance in tEVs (Fig. 5A,D), suggesting that, indeed, proteins are involved in the immobilization of EVs.

**Figure 5.**
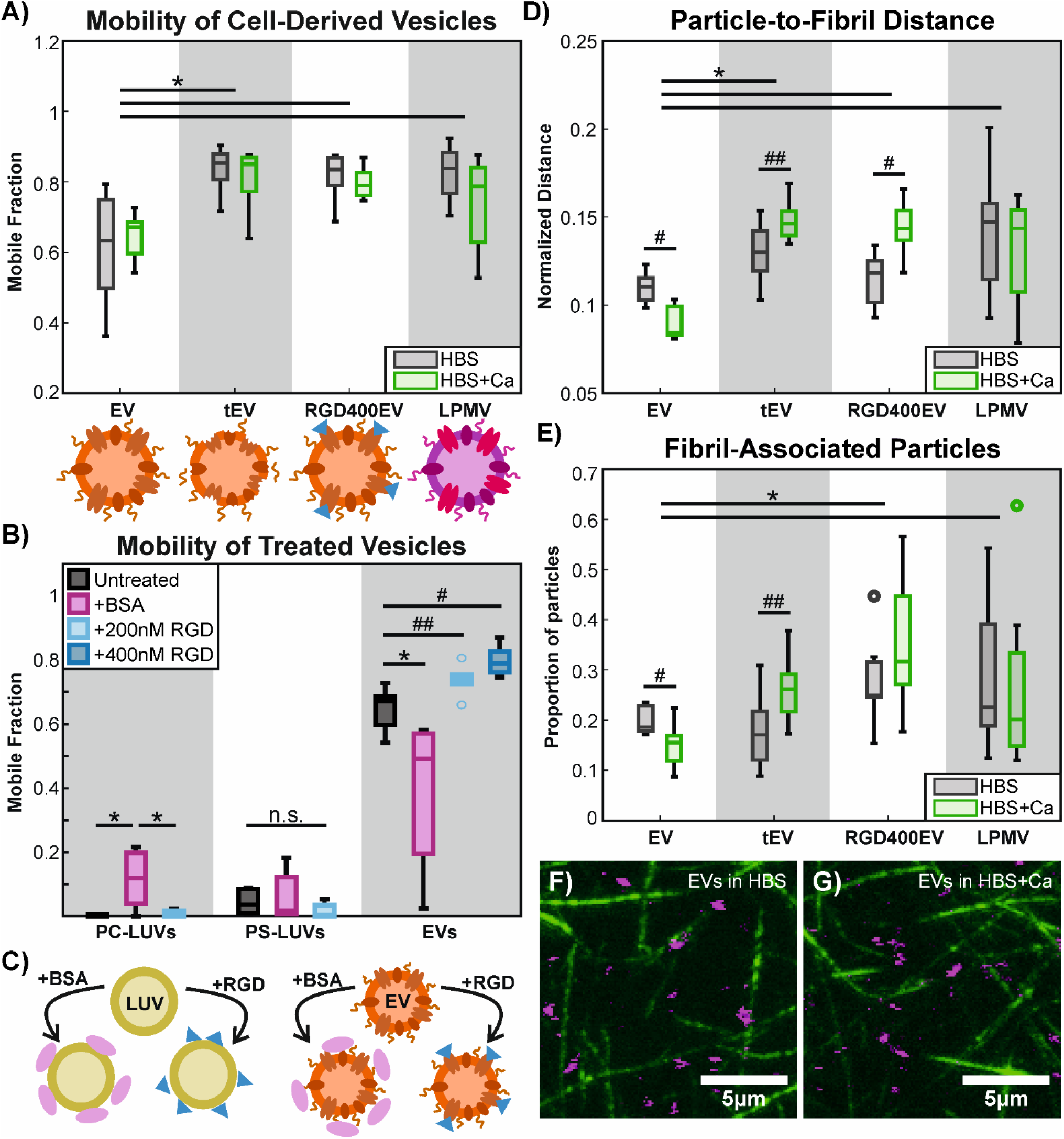
EV mobility in collagen I hydrogels is influenced by RGD-binding receptors. A) Mobile fractions of EVs diffusing in collagen I hydrogels are compared with those of trypsinized EVs (tEV), EVs treated with 400 nM cyclic RGD-peptide (RGD400EV), and LPMVs. The presence of calcium does not significantly affect the mobile fraction (p>0.05), but EVs are significantly less mobile than tEVs, pepEVs, and LPMVs according to a 2-way ANOVA with Tukey-Kramer post-hoc analysis, as indicated with * (p<0.01) with n=7 replicates. Cartoons below compare the composition and state of the particles. From left to right, an intact EV is compared to a trypsinized EV with its surface proteins digested, an EV treated with cyclic RGD peptides (blue triangles), and an LPMV with a membrane composition closer to that of whole cell plasma membrane. B) As extra controls, PC-LUVs and PS-LUVs were also treated with the RGD peptide to determine the effect of non-specifically adsorbed peptide on the particle surface. Treatment of EVs and LUVs with BSA served as a further negative control, as BSA should adsorb to particle surfaces without specifically inhibiting RGD-binding integrins. The RGD peptide does not improve the mobilities of PC- or PS-LUVs (p>0.05), but appears to increase EV mobility in a dose-dependent manner according to 2-sample T-tests when compared with untreated EVs, as indicated by # (p<0.01) and ## (p<0.05). BSA increases PC-LUV mobility, as determined by 1-way ANOVA across the PC-LUV conditions, as indicated by * (p<0.01). It has no effect on PS-LUVs (p>0.05) and decreases EV mobility (p<0.01), according to separate 1-way ANOVA tests. Data consist of n=6 replicates for LUVs and n=7 replicates for EVs. Data for EVs are the same from previous figures, but used for separate statistical comparisons. C) Cartoons depicting the effects of BSA and the RGD peptide when adsorbed on LUV and EV surfaces. Both BSA and the RGD peptide would adsorb nonspecifically on LUV surfaces, while the RGD peptide would bind specifically to receptors on EVs. D) Average particle-to-fibril distances of EVs, tEVs, pepEVs, and LPMVs, normalized by hydrogel mesh size. EVs are found significantly closer to collagen fibrils compared to tEVs, RGD400EVs, and LPMVs. Calcium further decreases the particle-to-fibril distance in EVs, but has the opposite effect in tEVs and RGD400EVs. LPMVs appear insensitive to the presence of calcium (p>0.05). E) Comparison of the proportion of fibril-associated particles, as determined by proximity. There are significantly less EVs colocalizing with collagen fibrils compared to RGD400EVs and LPMVs. Calcium appears to further decrease colocalization in EVs, but the opposite is true for tEVs. RGD400EVs and LPMVs are unaffected by calcium. For D and E, significant differences between particle types are determined with 2-way ANOVA with Tukey-Kramer post-hoc analysis, as indicated by * (p<0.05). Significant differences due to the presence of calcium within the same particle condition are determined with 2-way T-tests, as indicated by # (p<0.01) and ## (p<0.05). RGD400EVs and LPMVs were unaffected by calcium (p>0.05). Data consist of n=8 replicates across all conditions for analysis of particle-to-fibril distance and fibril association. F,G) Composite confocal images of collagen fibrils (green; image obtained in reflection mode) and fluorescently labelled EVs (magenta) in calcium-free (F) and calcium-containing (G) buffer. Images consist of a projection in the Z-axis over 10 slices with 0.8 μm spacing. Full-sized images of the 40×40 µm images from which these images were cropped can be found in Suppl. Fig. S10.

We next wanted to further narrow down the identity of the protein or proteins involved in EV immobilization. A previous study found that microvesicles derived from myofibroblasts specifically bound collagen I *via* integrin complexes containing the ITGB1 subunit.^39^ To see if this was the case here, we treated our EVs with a cyclic argynylglycylaspartic acid (RGD) peptide as a competitive binder and inhibitor of RGD motif-binding integrins (RGD400EVs).^87–90^ From our Western blot analysis of protein expression in EVs, we know that ITGB1 is highly enriched in the EV membrane. ITGB1 forms numerous ECM-binding integrin complexes with a variety of α-subunits, several of which are known to bind to RGD motifs found in fibronectin, vitronectin, and fibrinogen.^91,92^ Importantly, RGD motifs can be found in collagen I, but are only exposed and available for binding upon full or partial denaturation of the collagen molecule.^93,94^ Upon treatment of EVs with the RGD peptide, we observe an increase in overall particle mobility (Fig. 4A; see distribution of log_10_ diffusion coefficients in Suppl. Fig. S8). This supports the existence of non-fibrillar, unincorporated or denatured collagen in our hydrogels, similar to what has previously been reported.^95^ To be sure that this was due to the specific inhibition of RGD-binding integrins, we also tested the effect of the peptide when it nonspecifically adsorbed onto the surface of PCand PS-LUVs (Fig. 5B). The RGD peptide does not improve the mobilities of the PCand PS-LUVs, but appears to have a dose-dependent effect on EVs. This is most likely not due to electrostatic effects, as the surface charge of EVs is unchanged by treatment with the peptide (Suppl. Fig. S9A). DLS analyses of size distributions show that the peptide may induce small amounts of aggregation at 400 nM (Suppl. Fig. S9C), but this would have the opposite effect of making EVs less mobile. Furthermore, the mesh size of the collagen gels remains sufficiently larger than the size of the aggregates, such that steric interactions should be minimal. Thus, we conclude that RGD-binding integrins modulate EV mobility in collagen I hydrogels. To further validate this, we also treated our LUVs and EVs with bovine serum albumen (BSA) as a final negative control to see the effect of a ‘blocking’ agent that would not specifically bind to or prevent engagement between integrins and matrix collagen. The BSA improved the mobility of the PC-LUVs, but not the PS-LUVs. The decrease in EV mobility when treated with BSA can be attributed to interactions between the surface of the adsorbed BSA layer and the surrounding matrix environment, as the mobile fraction is intermediate between untreated EVs and the BSA-treated LUVs.

Altogether, this further supports our hypothesis that the peptide is specifically inhibiting integrin interactions with matrix collagen. Moreover, the magnitude of the increase in EV mobility upon treatment with 400 nM RGD peptide suggests that EV diffusion is more heavily influenced by interactions with non-fibrillar collagen *via* RGD-binding integrins than by those with fibrillar collagen. The sum of the mobile fraction (≥0.8) with the proportion of fibril-associated particles (0.2∼0.45) of RGD400EVs in both calcium-free and calcium-containing media exceeds 1, meaning these two populations of particles likely overlap and that some of these fibril-associated particles might not be fully immobilized. This suggests a degree of transience in these interactions with fibrillar collagen, allowing particles to bind and unbind as they diffuse along the collagen fibrils.

While their RGD-binding receptors would be inhibited, EVs treated with RGD peptide may still possess other uninhibited receptors, such as those that bind the GFOGER sequence on fibrillar collagen I.^70,92,96,97^ These receptors may require divalent cations other than calcium,^96,97^ however, which would explain the apparently weak interactions between RGD-treated EVs and the collagen fibrils. Further analysis involving targeted knock-down or inhibition of specific αand βintegrin subunits would be required to identify exactly which integrin complexes are involved.

The role of calcium in particle interactions with collagen appears to be context-specific. In synthetic PS-LUVs, negative surface charge appears to increase particle mobility in calcium-free buffer, but calcium ions appear to cause ‘bridging’ interactions between phosphatidylserines and matrix collagen that cause nearly full immobilization. In tEVs, the digestion of surface proteins by trypsin tends to target lysine and arginine residues, both of which are positively charged.^98^ Despite the overall surface charge not changing significantly after trypsinization (Suppl. Fig. S9), the increase in fibril association in the presence of calcium may also be due to electrostatic ‘bridging’ between exposed, charged protein fragments and collagen fibrils. The particle-to-fibril distances of tEVs and RGD400EVs increases in response to calcium, similar to PEG-LUVs and opposite to untreated EVs. This increase in particle-to-fibril distance may therefore be a consequence of non-specific charge interactions on membranes, while the slight decrease, as seen in untreated EVs would be due to specific receptor-ligand interactions, as discussed above. LPMVs do not appear to be affected by calcium, possibly due to alteration of surface proteins by the vesiculation agent or more likely because of overall differences in membrane composition.

The existence and localization of non-fibrillar collagen within the fibrillar mesh network of our hydrogels is difficult to definitively prove with imaging. Contrast in confocal reflectance microscopy is generated from backscattered light off of materials with sufficiently different refractive indeces.^99^ Collagen fibrils, due to their dense aggregation and regular alignment of molecules have a strong enough optical density to show up in images. While this imaging technique is not specific for fibrillar collagen like second harmonic generation,^100^ it is unable to pick out materials whose optical qualities are too similar to the aqueous background. This automatically excludes molecules in solution, as well as any polymeric materials that hold onto substantial amounts of water, such as hyaluronic acid and other glycosaminoglycans and proteoglycans.^73,101^ We can therefore be fairly confident that our images with confocal reflectance primarily represent non-denatured fibrillar collagen I. Visualization of non-fibrillar collagen would not be possible without labelling and possibly altering the molecular structure of the collagen. Our hydrogels were also too dilute for Raman microscopy with the Raman signal too weak for differentiating between fibrillar and non-fibrillar structures. It is, thus, only possible to infer the existence of non-fibrillar collagen in our hydrogels from our data.

### Implications of EV-matrix interactions

In order to systematically and quantitatively study EV-matrix interactions, we have used purified collagen I hydrogels to model the tissue microenvironment. Future work, however, might involve other structural ECM proteins, glycoaminoglycans, or more complex mixtures and materials, such as interpenetrating networks of hyaluronic acid, fibronectin, or even decellularized ECM. As we have shown, structural ECM components may not be the only molecular species that influence particle diffusion, but colloidal and soluble materials may also play an important role in regulating the diffusion of EVs. Furthermore, we expect the continued development of intravital imaging to yield important data that can link EV diffusion in living tissues with systemic distribution patterns *in vivo*.^34,87,102,103^ Working gradually and systematically towards model environments of greater complexity will allow teasing apart of individual interactions between EVs and ECM components and determination of how they impact EV diffusion as a whole in real tissues.

Our results showing that cyclic RGD peptides can be bound by EVs and that this appears to increase EV mobility within a reconstituted ECM environment has important implications for the use of RGD peptide-based drugs^87–89^. Because EVs are found nearly ubiquitously in biological fluids^1,3,33^, they have the potential to act as sinks for therapeutic substances, taking up drug molecules meant to reach target cell populations and lowering overall treatment efficacy. Moreover, the use of RGD peptides as integrin inhibitors to treat cancers^87–89^ may have the unintended effect of allowing cancer cell-derived EVs to become more mobile, increasing their range and ability to reach cells in distant tissues. Increasing evidence points towards EVs having the ability to prime tissue environments for metastatic invasion by reprogramming stromal cells or modifying the local ECM environment^2,14,16,19,20,40,55^. While inhibition of specific integrin complexes may prevent settling and trapping of EVs in certain tissues,^19,40^ the presence of other integrin complexes may simply allow them to settle in other tissue environments. It does not seem feasible with current technology to attempt to inhibit all integrin species in EVs to prevent them from settling anywhere. Research on integrin-inhibiting therapeutics would therefore benefit from *in vivo* studies involving analysis of circulating EVs and their ultimate distribution patterns in different tissues.^34^ Our results suggest, however, that it is possible to control diffusion and infiltration of EV-like particles to deliver therapeutic materials to cells in specific tissue microenvironments. Tailoring surface properties of nanoscale vehicles for therapeutics to interact with ECM materials could allow for more precise delivery mechanisms.

## Conclusions

Tissues consist of complex microenvironments and the diffusion of particles within them is governed by different kinds of non-specific physical and electrostatic,^74^ as well as biochemical interactions. EVs contain all the necessary machinery for interfacing with ECM materials and it is the expression of different kinds of surface receptors that determines what kind of interactions are possible. While we can emulate the basic biophysical properties of EVs with synthetic lipid vesicles, their interactions with ECM components and overall diffusive behaviour are controlled by more specific biochemical factors.

Compared to EVs, synthetic lipid LUVs tend to be immobilized in collagen I hydrogels unless a buffer layer of adsorbed colloidal proteins or PEG as a surface crowder exists to prevent adhesion. We also compared the behaviour of EVs to LPMVs, membrane structures of similar size and shape to EVs but with a composition that should more closely reflect that of whole cell plasma membrane. Despite coming from the same source cell type, EVs and LPMVs differ not only in composition and biophysical state, but also in terms of their interactions with collagen I and resulting diffusive behaviour. Finally, by using different strategies to modify EV interactions with collagen I, we were able to identify RGD-binding integrins as major determinants of EV mobility.

Trypsinization and treatment with a cyclic RGD peptide allowed us to narrow down the membrane component responsible for immobilization in collagen I hydrogels; first to a protein, then to an RGD-binding integrin complex. The ability of RGD peptides to increase EV mobility has important implications for the use of RGD peptide-based drugs, especially for treating cancer, but also shows that the diffusion and distribution of nanoparticles can be controlled by altering surface interactions. With more precise control over these interactions, nanoparticles could be designed to infiltrate and target specific tissues or be retained in others.

In summary, our key findings are that (i) EVs membranes have distinct compositions and biophysical properties different from whole cell membranes; (ii) EV diffusion in collagen I hydrogels is governed by surface interactions, i.e. charge and receptor-ligand binding; (iii) EVs appear to interact with non-fibrillar collagen *via* RGD-binding integrins; and (iv) EV mobility can be enhanced by inhibiting RGD-binding integrins with synthetic RGD peptides

## Materials and Methods

### Cell Culture

MDA-MB-231 breast cancer cells were obtained from the American Type Culture Collection and cultured in 100mm-diameter Petri dishes at 37 °C under 5% CO_2_. The complete culture medium consisted of low-glucose Dulbecco’s Modified Eagle’s Medium (DMEM; Sigma-Aldrich, USA) supplemented with 10% fetal bovine serum (FBS; Thermo Fisher Scientific, USA) and 1% penicillin-streptomycin (Thermo Fisher Scientific, USA). Cells were passaged every 3-4 days at 80-90% confluency as follows: old medium was removed and cells were washed twice with phosphate buffered saline (PBS; 137 mM NaCl, 2.7 mM KCl, 10 mM Na_2_HPO_4_, 1.8 mM KH_2_PO_4_). Next 2 mL trypsin/EDTA solution (PAN-Biotech, Germany) was added and the cells returned to the incubator for 5 minutes to allow detachment. The trypsin was quenched with an addition of 2 mL complete culture medium before being collected and centrifuged at 200 ×g for 10 minutes. After pelleting, the supernatant was disposed and the pellet was resuspended in fresh complete culture medium. Cells were plated in Nunc cell culture-treated Petri dishes (Thermo Fisher Scientific, USA) with 7-8 mL of culture medium at an approximate split ratio of 1 to 3 or 4.

### Buffers

In order to study the effects of calcium on membrane interactions with collagen, we could not use PBS due to the tendency for calcium to crash out of solution as calcium phosphate. Instead, we opted to use a calcium-containing buffer previously described for use in generating giant plasma membrane vesicles (GPMVs)^48^, referred to as GPMV buffer and consisting of 150 mM NaCl, 10mM HEPES, and 2 mM CaCl_2_; pH 7.4. In this manuscript, we refer to this buffer as HBS+Ca to contrast it with our calcium-free buffer, which is a custom HEPES buffered saline (HBS) mixture consisting of 150 mM NaCl and 16mM HEPES, pH 7.4. In this buffer, the calcium was removed from the original HBS+Ca recipe and the amount of HEPES was increased to compensate for the loss in osmolality. The osmolality of both buffers was determined to be 303 mOsm/kg with a Gonotech freezing point osmometer (Gonotech, Germany). The amount of calcium in HBS+Ca is representative of extracellular calcium concentrations in tissues^104^.

### Generation and purification of EVs

To obtain enough EVs for experiments, 10-12 plates of cells were cultured normally to 80-90% confluency. To avoid contamination with bovine vesicles, cells were switched to a serum-free medium. First, plates of cells were washed 3 times with PBS before the medium was replaced with 7mL low-glucose DMEM supplemented with 1% penicillin-streptomycin. Cells were incubated for 3 days in serum-free conditions to generate EVs. Serum-starvation should also have enhanced the number of EVs generated^10^. Conditioned media were collected and pooled together, then centrifuged at 400 ×g for 10 minutes to pellet dead cells which may have been lifted off the plate. The supernatant was retained and centrifuged at 2000 ×g to remove remaining cell debris before being concentrated using 100kDa Amicon Ultra-15 centrifugal filters (Merck Millipore, USA), centrifuged at 3400 ×g to a final volume of 1 mL. The concentrated conditioned medium was then incubated for 10 minutes with 1 μL 2.5 mg·mL^-1^ 1,1’-Dilinoleyl-3,3,3’,3’-Tetramethylindocarbocyanine, 4-Chlorobenzenesulfonate (FAST DiI; Fischer Scientific, USA) dissolved in ethanol or 1 μL 2.5 mg/mL N,N-Dimethyl-6-dodecanoyl-2-naphthylamine (LAURDAN; Thermo Fisher, USA) dissolved in dimethyl sulfoxide to label the EVs. The sample was then run through a homemade size exclusion chromatography column made with a 10 mL syringe with the plunger removed, packed with Sepharose CL-4B base matrix (Sigma-Aldrich, USA), and eluted with gravity flow.

To equilibrate the Sepharose, approximately 15 mL of suspended Sepharose matrix was allowed to settle in a 50mL conical tube for 2 hours and the liquid medium was removed and replaced with fresh HBS. This was repeated 5 times to wash the Sepharose beads. A syringe was prepared by stopping it with an end cap and a Whatman polycarbonate membrane filter with 10μm pore size (Sigma-Aldrich, USA) was cut to fit in the bottom to prevent the Sepharose beads from coming out. After the final wash, the Sepharose beads were suspended in HBS and loaded into the prepared syringe and left overnight to pack. Two column volumes (20 mL) of HBS were run through the column before the sample was added. Up to 20 fractions of approximately 500 μL each were collected with 1mL additions of HBS at a time. Fractions 7-10 were found to be enriched in particles 100-400 nm in diameter, as determined with dynamic light scattering (see SF. 1). These were pooled together and re-concentrated using centrifugal filter tubes with a molecular weight cutoff of 100 kDa. To obtain vesicles in a calcium-containing buffer, EVs were concentrated and resuspended in HBS+Ca before re-concentrating. This was repeated 5 times to replace the medium.

### Generation of GPMVs and extrusion of LPMVs

GPMVs were generated according to a previously reported protocol^48^. Briefly, 10-12 plates of 80-90% confluent MDAMB-231 cells were washed twice with PBS, then once with HBS+Ca (referred to as GPMV buffer in the original protocol). A stock solution was made of 1M N-ethylmaleimide (NEM; Sigma-Aldrich, USA) dissolved in distilled water. This was stored at -20 °C and thawed before use. The buffer in each plate was then replaced with 2 mL vesiculation buffer, consisting of HBS+Ca plus 2 μL NEM stock per 1 mL of HBS+Ca buffer. Plates were left to incubate at 35 °C for 1 hour to allow for vesiculation. GPMVs were then collected by gentle pipetting, avoiding uplift of the cells. The vesiculated material was centrifuged at 100 ×g for 10 minutes to remove cell debris, then at 20000 ×g for 1 hour to pellet the GPMVs. The supernatant was removed and the pellet resuspended in 1mL HBS+Ca. This material was then incubated with 1 μL FAST DiI or LAURDAN, similarly to EVs for 10 minutes to label the membranes. The labelled membranes were extruded with an Avanti handheld extruder fitted on a heating block (Avanti Polar Lipids, USA) set on a hot plate at 37 °C. Extrusion was done in 2 steps, first 21 passes through a Whatman Nuclepore 400 nm-pore size track-etched membrane filter, then 21 passes through a 200 nm-pore size filter (Sigma-Aldrich, USA). If done at room temperature, the membrane material tended to clog the pores of the filter. At 37 °C, the membrane was in the fluid state, allowing greater ease in extrusion and ensuring the inclusion of lipids that would otherwise be in the gel state at room temperature. The resulting LPMVs were concentrated and washed the same way EVs were.

### CryoSEM of EVs and LPMVs

EV and LPMV suspensions were concentrated approximately tenfold with centrifugal filter tubes with a molecular weight cutoff of 100 kDa. Sample volumes of around 14 µL each were sandwiched between two type B gold-coated freezing discs (BALTIC Preparation, Germany) and high pressure frozen with a Leica EM HPM100 High Pressure Freezer (Leica Microsystems, Austria). Samples were mounted under liquid nitrogen onto a cryo-sample holder in a Leica EM VCM Mounting Station (Leica Microsystems, Austira), then transferred using a VCT500 shuttle (Leica Microsystems, Austria) to a Leica EM ACE600 system (Leica Microsystems, Austria) for freeze-fracturing and sputter coating. Particles embedded in the surrounding frozen medium were exposed by freeze-fracturing and an 8 nm-thick platinum film was applied at -160°C. Samples were transferred to a Quattro Environmental Scanning Electron Microscope (Thermo Fisher Scientific, USA) under high vacuum at a pressure of 3.08×10^−7^Torr. An Everhart-Thornley detector was used with an acceleration voltage of 5.00 kV.

### Production of LUVs

All lipids were purchased from Avanti Polar Lipids (USA) and were dissolved in chloroform. LUVs were produced from lipid mixtures consisting of 4 mM DOPC, 4 mM DOPC +1 mol% DSPE-mPEG5K, or 4 mM 4:1 DOPC/DOPS with 0.5 mol% DiI (Fisher Scientific, USA) as a fluorescent label. First, 20 μL of lipid was spread and dried in a glass vial under vacuum for 1 hour. Next, the lipid was rehydrated with 1 mL buffer, either HBS or HBS+Ca, then vortexed for 5 minutes to form multilamellar structures. These were then extruded 21 passes with an Avanti handheld extruder fitted with a 200 nm pore size Whatman Nuclepore track-etched membrane filter.

### DLS characterization of size and surface charge

Dynamic light scattering (DLS) was used as a first pass at characterizing the size distribution of EVs and LPMVs. It was also used to ensure quality and uniformity of extruded synthetic LUVs. Samples of particles were loaded into disposable folded capillary tubes (DTS1070; Malvern Panalytical, UK) and measured with a Malvern Instruments Nano-ZS Zetasizer equipped with a 632.8 nm 4 mW HeNe laser (Malvern Panalytical, UK). Size distributions were obtained at a scattering angle of 173° before determination of zeta potential. The high salt buffer conditions likely resulted in electrostatic screening, so the zeta potential here is presented as a relative measure of surface charge.

### Western blot characterization of protein expression

Ten plates’ worth of EVs and LPMVs, along with the nonEV fractions obtained with SEC (fractions 1-6, 11-20) were concentrated in centrifugal filters to a final volume of approximately 200 μL. These were then lysed using 20 μL radioimmunoprecipitation assay (RIPA) buffer at 10× concentration (1.5 M NaCl, 10% Nonidet-P40, 5% sodium deoxycholate, 1% sodium dodecyl sulfate [SDS], 500 mM Tris). Lysates were stored at -80 °C until use. Whole cell lysates were obtained by adding 2 mL 1× RIPA buffer to one plate of 90% confluent MDA-MB-231 cells and scraping and pipetting to dislodge the cell material. A Bradford assay was used to determine the amount of whole protein in the lysates. Briefly, 10 μL of each sample was diluted to a final volume of 100μL in a 96-well plate with PBS. Samples were then serially diluted by adding 100 μL PBS, pipetting up and down to mix, and then transferring 100μL of the mixed sample to a new well. This was repeated to obtain a dilution series. Each well then received 100 μL of Bradford reagent (Thermo Fisher, USA). The plate was gently shaken to mix the well contents and the absorbance at 595 nm was measured with a Biotek Cytation 5 Microplate Reader (Agilent, USA). The absorbances of the diluted samples were used to form dilution curves and the slopes of the linear regions of each curve were compared to determine the necessary dilution factors for each lysate that would allow the curves to overlap. The original samples were then diluted accordingly with distilled water to normalize for whole protein content.

For polyacrylamide gel electrophoresis, Laemmli Buffer (Bio-Rad, USA) at 2× concentration was supplemented with βmercaptoethanol (Sigma-Aldrich, USA) according to the manufacturer’s instructions, then added 1:1 to samples before heating to 95 °C for 5 minutes. Once cooled, samples were vortexed to homogenize them, then loaded in 25 μL volumes onto a homemade 10-well 10% acrylamide stacked SDS-PAGE gel with a 4% acrylamide stacking layer. Progression of the separation of molecular weights was visualized with a Spectra Multicolor Broad Range Protein Ladder (Thermo Fisher, USA). Electrophoresis was run at 100 V for 1 hour, then at 150 V for 45 minutes in homemade Tris-glycine-SDS running buffer (25 mM Tris, 192 mM glycine, 0.1% SDS) using a Bio-Rad MiniPROTEAN Tetra cell, tank, and power supply (Bio-Rad, USA). The proteins were then transferred to a polyvinylidene fluoride (PVDF) membrane (Bio-Rad, USA) with the Bio-Rad wet blotting cell at 200 V for 90 minutes in Tris-glycine transfer buffer (25 mM Tris, 192 mM glycine, 20% v/v methanol). After transfer, the PVDF membrane was equilibrated in Tris-buffered saline (20 mM Tris, 150 mM NaCl) with 0.1% Tween-20 (TBST) before being blocked overnight at 4 °C or for 1 hour at room temperature with gentle orbital shaking in 2.5% w/v bovine serum albumin (BSA; Sigma-Aldrich, USA) dissolved in TBST. The proteins, ALIX, CD63, and ITGB1 were probed in that order as follows: after blocking, the blot was rinsed with TBST, then allowed to incubate overnight at 4°C with primary antibody, diluted 1:1000 in 1% BSA dissolved in TBST. The blot was then rinsed three times with TBST, then incubated with the appropriate horseradish peroxidase-conjugated secondary antibody, diluted 1:1000 in 1% BSA dissolved in TBST for an hour at room temperature. Blots were, again, rinsed three times with TBST before 750 μL each of two Pierce ECL Western Blotting Substrate solutions (Thermo Fisher, USA) were added. Blots were shaken and allowed to incubate at room temperature for 5 minutes before imaging with a G:Box Chemi XX6 gel documentation system (SynGene, UK) under chemiluminescence mode. After imaging, blots were rinsed with TBST, then stripped to allow subsequent probing of other proteins. A mild-strength acidic stripping buffer was used, consisting of 200 mM glycine, 0.1% SDS, 1% Tween-20, pH 2.2. A small amount of stripping buffer was first added to blots to lower the pH, then discarded before enough was added to cover the blot. The blot was then left to strip for 20 minutes at room temperature with gentle rocking. The buffer was next discarded, and the blot was stripped a second time as before. To neutralize the blot pH, the blot was rinsed three times with TBST, then blocked with 2.5% BSA in TBST for the next round of antibody treatment.

The following primary antibodies were purchased from Thermo Fisher (USA) and used for Western blot analysis: ALIX recombinant rabbit monoclonal antibody (JM85-31), CD63 mouse IgG1 monoclonal antibody (Ts63), ITGB1 mouse IgG1 monoclonal antibody (3B6). The following secondary antibodies were used with the appropriate primary antibody, according to source species selectivity: Goat anti-mouse IgG (H+L) HRP-conjugated secondary antibody (Thermo Fisher, USA) and goat anti-rabbit HRP-linked IgG secondary antibody (Cell Signaling Technology, USA). Full, uncropped versions of blots and densitrometric analysis over 6 replicates can be found in the Supplemental Materials, Fig. S4.

### LAURDAN fluorescence spectroscopy and phasor analysis

EVs and LPMVs labelled with 0.5 mol% LAURDAN were loaded in a quartz crystal cuvette (Hellma, Germany) and their fluorescence spectra were measured with 360 nm excitation using a FluoroMax Plus Spectrofluorometer (Horiba, Japan). The spectral information were then converted to phasor coordinates, as previously reported^61–63^ with the following equations:

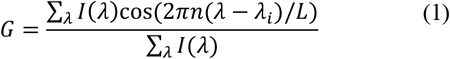

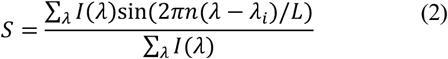

Where *I(λ)* is the measured fluorescence intensity at wavelength, λ; *n* is the harmonic number (here taken to be 1); *λ*_*i*_ is the initial or shortest wavelength measured; and *L* is the length of the spectrum. Once converted to phasor coordinates, the data points were plotted on a Cartesian plane, where G (Eq. 1) was the x-coordinate and S (Eq. 2) was the y-coordinate.

We also measured the fluorescence spectra of multilamellar membranes produced from the following lipid mixtures, dissolved at 4 mM overall lipid concentration with 0.5 mol% LAURDAN in chloroform: pure DOPC, 70% DOPC + 30% cholesterol (DOPC+Chol), and 13% DOPC + 44% dipalmitoylphosphatidylcholine (DPPC) + 43% cholesterol (Ternary L_O_). Cholesterol was purchased from Sigma-Aldrich (USA). All other lipids were purchased from Avanti Polar Lipids (USA). The multilamellar membranes were produced by drying 10 μL of the lipid mixtures in glass vials under vacuum for 1 hour before rehydrating in 500 μL buffer (either HBS or HBS+Ca). Phasor analysis of the synthetic membranes produced a linear trajectory showing increasing packing, membrane dehydration, and membrane order from DOPC to the ternary L_O_ mixture. Alignment with this trajectory in EVs and LPMVs suggests a simple biophysical state that can be replicated with minimal lipid membrane systems. Information on lipid phase separation cannot be gleaned from this analysis, as the signals from LAURDAN molecules in different phases would sum as a linear combination, resulting in a mean spectrum resembling that of a single phase of intermediate membrane order.

### EV treatments to inhibit integrin function

To determine if proteins were responsible for EV immobilization in collagen I hydrogels, EVs were trypsinized with TrypLE™ Express Enzyme (Thermo Fisher, USA). EV suspensions were first concentrated down to approximately 100 μL with centrifugal filter tubes with a molecular weight cutoff of 100 kDa. Next, 200 μL of trypsin solution was added directly to the EV suspension in the centrifugal filter tubes. This was allowed to incubate at 37 °C for 10 minutes before being diluted with HBS or HBS+Ca, depending on the desired final buffer condition to fill the filter tube (approximately 800 μL). The sample was then centrifuged at 3400 ×g to concentrate it down to 100 μL before being washed a further 4 times in this manner to remove the trypsin. EV size distributions and surface charge before and after trypsinization were measured with DLS (SF.6). To specifically inhibit RGD motif-binding integrins, EVs were treated with a cyclic RGD peptide with the following amino acid sequence: GGGGCRGDSPC (Peptide 2.0, USA). A 12mM stock solution was prepared in distilled water, then aliquoted and diluted 1:1 with 2× concentration HBS or HBS+Ca. This was then added to EV suspensions to a final concentration of 120nM peptide. Previous studies on the use of cyclic RGD peptide-based drugs has shown that receptor affinity varies with integrin subtype, with IC_50_ values spanning several orders of magnitude, but generally in the nano-molar range^88,90^.

As negative controls, EVs and LUVs were treated with 200 nM RGD peptide or 0.1 mg·mL^-1^ bovine serum albumen (Sigma-Aldrich, USA) in appropriate buffers and incubated for 10 minutes at 37 °C prior to pipetting into collagen gels.

### Formation of collagen I hydrogels

To form 50 μL hydrogels, 12.5 μL of a 6mg/mL stock solution of solubilized collagen I from bovine skin (Sigma-Aldrich, USA) was pipetted into a well of a 96-well plate. The plate and the stock were kept on ice to prevent premature gelation. The pH of the collagen was corrected to approximately 7 with 1 μL 1 M NaOH, then diluted 1:1 with 12.5 μL 2× concentrated buffer (HBS or HBS+Ca). The solution was mixed by pipetting up and down before being further diluted 1:1 with 24 μL 1× concentrated buffer, resulting in a final dilution factor of 1:4 and an in-gel collagen I concentration of 1.5 mg/mL. To maintain humidity, surrounding wells of the 96-well plate were filled with distilled water. The whole plate was then sealed with parafilm and placed in an incubator at 37 °C to allow gelation overnight.

### Single particle tracking and analysis of mobility

To maintain the same approximate particle concentration that is added to gels across sample preparations, labelled vesicles were imaged with a pco.Edge sCMOS camera (PCO AG, Germany) mounted on a Zeiss AXIO Observer.D1 microscope equipped with a 63× 1.2NA water immersion C-Apochromat objective (Carl Zeiss, Germany) in epifluorescence mode. The number of particles in solution was counted with a particle detection function in FIJI and the particle suspensions were diluted accordingly. Particles were introduced to hydrogels by pipetting 10 μL of suspension onto collagen gels and allowing them to incubate at 37 °C for minimum 3 hours to allow particles to diffuse throughout.

Image sequences of diffusing particles in 80×80 μm regions in hydrogels were collected as above with a pco.Edge sCMOS camera at a frame rate of 20fps with ∼45 ms exposure. The particle tracking plugin developed for FIJI by Sbalzarini and Koumoutsakos^44^ was used to identify particles and determine their diffusion coefficients. For analysis of mobility, the base-10 logarithms of the diffusion coefficients were determined and visualized with histograms normalized such that the bins sum up to 1 (SF.3 and 5). To determine the mobile fraction, the bins of the histogram above a log_10_ diffusion coefficient value of -13 were summed. In a previous publication, we were able to define a threshold based on the noise floor of our imaging system for determining whether a particle was mobile^74^. Here, we were unable to define such a threshold because the particle size was not as well-defined, with a relatively high degree of polydispersity especially in EVs. As such, we chose -13 as a value that appears to clearly separate the bimodal peaks of the distributions we obtained, and that corresponds to a particle that is definitely ‘mobile’.

### Confocal microscopy of collagen gels

Collagen fibrils were imaged in confocal reflection mode^72,73^ with a Leica SP8 FALCON microscope equipped with a 63× 1.2 NA water immersion objective (Leica, Germany) with 488 nm argon laser illumination. Z-stacks consisting of 30 images were obtained with 0.75 μm spacing to get more data. In parallel, DiI-labelled particles were imaged with 561 nm diode laser excitation.

Hydrogel mesh size was determined according to a previously described procedure with slight modification^72^ (Fig. 3). Our confocal reflectance images contained scanner artifacts that we removed with a bandpass filter. This unfortunately introduced a spreading effect on the fibril pixels, so we binarized and skeletonized the images with FIJI to obtain the central axes of the fibrils. The exact width of the fibrils themselves would not be determinable with confocal microscopy, as they appear to be below the optical diffraction limit. As such, skeletonization does not remove any important information and allows mesh size, as well as particle-to-fibril distance to be defined by the central axes of fibrils, which we believe to be more accurate. Mesh size was then determined by counting up the number and sizes of spaces in the xand y-directions of the image between fibril pixels. The distribution of the sizes of these spaces fall into exponential distributions, where the mean value is defined by the exponent. Distributions were thus fit to exponential functions using the MATLAB curve fitting toolbox and the mean values were determined and converted from pixel values to lengths in micrometers. Thus, mesh size corresponds to the average length between fibrils.

To determine particle-to-fibril distance, fluorescence images of particles were binarized and the coordinates of the geometric centres of the particles were determined with FIJI. These coordinates could then be cross-referenced with the skeletonized images of the fibrils to determine the distance to the closest fibril. The Python implementation of the KD-Tree nearest neighbour search algorithm^86^ was used to determine these distances for each particle. We noticed that particle-to-fibril distance had a strong dependence on mesh size (see Suppl. Fig. S7), which tended to vary between samples. To eliminate this effect, we normalized particle-to-fibril distance to the mesh size, as determined above. We then defined a fibril-associated particle to be a particle that is colocalized or directly adjacent and touching a fibril. The noise floor for this would be approximately half the apparent width of the fibrils in our images (2-3 pixels) plus the apparent radius of the particles (also 2-3 pixels). The threshold distance for a fibril-associated particle was thus determined to be 5 pixels, corresponding to approximately 500 nm. The proportion of fibril-associated particles was then determined to be the proportion of particles whose particle-to-fibril distance fell below 500 nm.

### Statistics and data presentation

Boxplots show the median value of the dataset (middle line), first and third quartiles (box limits), and the maximum extent of non-outlier data (whiskers). Averaged histograms show mean value with error bars showing standard deviation. Statistical significance was determined with N-way ANOVA (as indicated) with Tukey-Kramer post-hoc analysis for multiple comparisons at the significance levels indicated. Two-sample T-tests were also conducted on selected isolated paired data sets to double check differences in the event they were not identified with ANOVA. Unless otherwise indicated, all other comparisons were found to not be statistically significant (p>0.05). Statistics were computed with MATLAB. Multiple collections of EVs and LPMVs were conducted to obtain experimentally independent replicates. Individual replicates for determining mobile fraction, particle-to-fibril distance, and fibril association consist of 100-500 identified particles. The variability in identified particles is partially due to differences in particle mobility and also due to inhomogeneities within and across hydrogel samples.

## Acknowledgements

The authors would like to acknowledge H. Runge and S. Weichold at the Max Planck Institute of Colloids and Interfaces for their help with cryoSEM. N.W. Tam would like to acknowledge funding from the International Max Planck Research School on Multi-Scale Biosystems. A. Becker would like to acknowledge funding from the German Academic Exchange Service (DAAD) RISE research internship program. A. Cipitria would like to acknowledge funding from the DFG Emmy Noether grant (CI 203/2-1), from IKERBASQUE Basque Foundation for Science, and from the Spanish Ministry of Science and Innovation (MCIN/AEI/10.13039/501100011033/FEDER UE, through grant PID2021-123013OB-I00).

## Supporting Information

Supplementary figures, including additional vesicle and hydrogel characterization data

## Author Contributions

N.W. Tam, A. Becker, and A. Mangiarotti designed and conducted experiments. N.W. Tam and A. Becker wrote the manuscript. R. Dimova and A. Cipitria contributed to project supervision and editing of the manuscript.

## REFERENCES

(1) Hessvik, N. P.; Llorente, A. Current Knowledge on Exosome Biogenesis and Release. Cell. Mol. Life Sci. 2018, 75 (2), 193–208. 10.1007/s00018-017-2595-9.

(2) Becker, A.; Thakur, B. K.; Weiss, J. M.; Kim, H. S.; Peinado, H.; Lyden, D. Extracellular Vesicles in Cancer: Cell-to-Cell Mediators of Metastasis. Cancer Cell 2016, 30 (6), 836–848. 10.1016/j.ccell.2016.10.009.

(3) Choi, H.; Lee, D. S. Illuminating the Physiology of Extracellular Vesicles. Stem Cell Res. Ther. 2016, 7 (1), 55. 10.1186/s13287-016-0316-1.

(4) Margolis, L.; Sadovsky, Y. The Biology of Extracellular Vesicles: The Known Unknowns. PLOS Biol. 2019, 17 (7), e3000363. 10.1371/journal.pbio.3000363.

(5) van Niel, G.; D’Angelo, G.; Raposo, G. Shedding Light on the Cell Biology of Extracellular Vesicles. Nat. Rev. Mol. Cell Biol. 2018, 19 (4), 213–228. 10.1038/nrm.2017.125.

(6) Pollet, H.; Conrard, L.; Cloos, A.-S.; Tyteca, D. Plasma Membrane Lipid Domains as Platforms for Vesicle Biogenesis and Shedding? Biomolecules 2018, 8 (3), 94. 10.3390/biom8030094.

(7) Haraszti, R. A.; Didiot, M.-C.; Sapp, E.; Leszyk, J.; Shaffer, S. A.; Rockwell, H. E.; Gao, F.; Narain, N. R.; DiFiglia, M.; Kiebish, M. A.; Aronin, N.; Khvorova, A. High-resolution Proteomic and Lipidomic Analysis of Exosomes and Microvesicles from Different Cell Sources. J. Extracell. Vesicles 2016, 5 (1). 10.3402/jev.v5.32570.

(8) Bissig, C.; Gruenberg, J. ALIX and the Multivesicular Endosome: ALIX in Wonderland. Trends Cell Biol. 2014, 24 (1), 19–25. 10.1016/j.tcb.2013.10.009.

(9) Hoshino, D.; Kirkbride, K. C.; Costello, K.; Clark, E. S.; Sinha, S.; Grega-Larson, N.; Tyska, M. J.; Weaver, A. M. Exosome Secretion Is Enhanced by Invadopodia and Drives Invasive Behavior. Cell Rep. 2013, 5 (5), 1159–1168. 10.1016/j.celrep.2013.10.050.

(10) Zou, W.; Lai, M.; Zhang, Y.; Zheng, L.; Xing, Z.; Li, T.; Zou, Z.; Song, Q.; Zhao, X.; Xia, L.; Yang, J.; Liu, A.; Zhang, H.; Cui, Z.; Jiang, Y.; Bai, X. Exosome Release Is Regulated by MTORC1. Adv. Sci. 2019, 6 (3), 1801313. 10.1002/advs.201801313.

(11) Hedlund, M.; Stenqvist, A.-C.; Nagaeva, O.; Kjellberg, L.; Wulff, M.; Baranov, V.; Mincheva-Nilsson, L. Human Placenta Expresses and Secretes NKG2D Ligands via Exosomes That Down-Modulate the Cognate Receptor Expression: Evidence for Immunosuppressive Function. J. Immunol. 2009, 183 (1), 340–351. 10.4049/jimmunol.0803477.

(12) Raposo, G.; Nijman, H. W.; Stoorvogel, W.; Liejendekker, R.; Harding, C. V; Melief, C. J.; Geuze, H. J. B Lymphocytes Secrete Antigen-Presenting Vesicles. J. Exp. Med. 1996, 183 (3), 1161–1172. 10.1084/jem.183.3.1161.

(13) Eguchi, T.; Sogawa, C.; Okusha, Y.; Uchibe, K.; Iinuma, R.; Ono, K.; Nakano, K.; Murakami, J.; Itoh, M.; Arai, K.; Fujiwara, T.; Namba, Y.; Murata, Y.; Ohyama, K.; Shimomura, M.; Okamura, H.; Takigawa, M.; Nakatsura, T.; Kozaki, K. ichi; Okamoto, K.; Calderwood, S. K. Organoids with Cancer Stem Cell-like Properties Secrete Exosomes and HSP90 in a 3D Nanoenvironment. PLoS One 2018, 13 (2), e0191109. 10.1371/journal.pone.0191109.

(14) Ono, M.; Kosaka, N.; Tominaga, N.; Yoshioka, Y.; Takeshita, F.; Takahashi, R. -u.; Yoshida, M.; Tsuda, H.; Tamura, K.; Ochiya, T. Exosomes from Bone Marrow Mesenchymal Stem Cells Contain a MicroRNA That Promotes Dormancy in Metastatic Breast Cancer Cells. Sci. Signal. 2014, 7 (332), ra63–ra63. 10.1126/scisignal.2005231.

(15) Saadeldin, I. M.; Kim, S. J.; Choi, Y. Bin; Lee, B. C. Improvement of Cloned Embryos Development by Co-Culturing with Parthenotes: A Possible Role of Exosomes/Microvesicles for Embryos Paracrine Communication. Cell. Reprogram. 2014, 16 (3), 223–234. 10.1089/cell.2014.0003.

(16) Shinde, A.; Paez, J. S.; Libring, S.; Hopkins, K.; Solorio, L.; Wendt, M. K. Transglutaminase-2 Facilitates Extracellular Vesicle-Mediated Establishment of the Metastatic Niche. Oncogenesis 2020, 9 (2), 16. 10.1038/s41389-020-0204-5.

(17) Lewin, S.; Hunt, S.; Lambert, D. W. Extracellular Vesicles and the Extracellular Matrix: A New Paradigm or Old News? Biochem. Soc. Trans. 2020, 48 (5), 2335–2345. 10.1042/BST20200717.

(18) Gradilla, A.-C.; González, E.; Seijo, I.; Andrés, G.; Bischoff, M.; González-Mendez, L.; Sánchez, V.; Callejo, A.; Ibáñez, C.; Guerra, M.; Ortigão-Farias, J. R.; Sutherland, J. D.; González, M.; Barrio, R.; Falcón-Pérez, J. M.; Guerrero, I. Exosomes as Hedgehog Carriers in Cytoneme-Mediated Transport and Secretion. Nat. Commun. 2014, 5 (1), 5649. 10.1038/ncomms6649.

(19) Nogués, L.; Benito-Martin, A.; Hergueta-Redondo, M.; Peinado, H. The Influence of Tumour-Derived Extracellular Vesicles on Local and Distal Metastatic Dissemination. Mol. Aspects Med. 2018, 60, 15–26. 10.1016/j.mam.2017.11.012.

(20) Costa-Silva, B.; Aiello, N. M.; Ocean, A. J.; Singh, S.; Zhang, H.; Thakur, B. K.; Becker, A.; Hoshino, A.; Mark, M. T.; Molina, H.; Xiang, J.; Zhang, T.; Theilen, T.-M.; García-Santos, G.; Williams, C.; Ararso, Y.; Huang, Y.; Rodrigues, G.; Shen, T.-L.; Labori, K. J.; Lothe, I. M. B.; Kure, E. H.; Hernandez, J.; Doussot, A.; Ebbesen, S. H.; Grandgenett, P. M.; Hollingsworth, M. A.; Jain, M.; Mallya, K.; Batra, S. K.; Jarnagin, W. R.; Schwartz, R. E.; Matei, I.; Peinado, H.; Stanger, B. Z.; Bromberg, J.; Lyden, D. Pancreatic Cancer Exosomes Initiate Pre-Metastatic Niche Formation in the Liver. Nat. Cell Biol. 2015, 17 (6), 816–826. 10.1038/ncb3169.

(21) Bebelman, M. P.; Smit, M. J.; Pegtel, D. M.; Baglio, S. R. Biogenesis and Function of Extracellular Vesicles in Cancer. Pharmacol. Ther. 2018, 188, 1– 11. 10.1016/j.pharmthera.2018.02.013.

(22) Kosaka, N.; Yoshioka, Y.; Fujita, Y.; Ochiya, T. Versatile Roles of Extracellular Vesicles in Cancer. J. Clin. Invest. 2016, 126 (4), 1163–1172. 10.1172/JCI81130.

(23) Xu, R.; Rai, A.; Chen, M.; Suwakulsiri, W.; Greening, D. W.; Simpson, R. J. Extracellular Vesicles in Cancer — Implications for Future Improvements in Cancer Care. Nat. Rev. Clin. Oncol. 2018, 15 (10), 617–638. 10.1038/s41571-018-0036-9.

(24) Lee, Y.; Ni, J.; Beretov, J.; Wasinger, V. C.; Graham, P.; Li, Y. Recent Advances of Small Extracellular Vesicle Biomarkers in Breast Cancer Diagnosis and Prognosis. Mol. Cancer 2023, 22 (1), 33. 10.1186/s12943-023-01741-x.

(25) Arif, S.; Moulin, V. J. Extracellular Vesicles on the Move: Traversing the Complex Matrix of Tissues. Eur. J. Cell Biol. 2023, 102 (4), 151372. 10.1016/j.ejcb.2023.151372.

(26) Kooijmans, S. A. A.; Vader, P.; van Dommelen, S. M.; van Solinge, W. W.; Schiffelers, R. M. Exosome Mimetics: A Novel Class of Drug Delivery Systems. International Journal of Nanomedicine. Dove Press 2012, pp 1525–1541. 10.2147/IJN.S29661.

(27) Sabanovic, B.; Piva, F.; Cecati, M.; Giulietti, M. Promising Extracellular Vesicle-Based Vaccines against Viruses, Including SARS-CoV-2. Biology (Basel). 2021, 10 (2), 94. 10.3390/biology10020094.

(28) L. Arias, J.; Clares, B.; E. Morales, M.; Gallardo, V.; Ruiz, M. Lipid-Based Drug Delivery Systems for Cancer Treatment. Curr. Drug Targets 2011, 12 (8), 1151–1165. 10.2174/138945011795906570.

(29) Ozpolat, B.; Sood, A. K.; Lopez-Berestein, G. Liposomal SiRNA Nanocarriers for Cancer Therapy. Adv. Drug Deliv. Rev. 2014, 66, 110–116. 10.1016/j.addr.2013.12.008.

(30) Das, C. K.; Jena, B. C.; Banerjee, I.; Das, S.; Parekh, A.; Bhutia, S. K.; Mandal, M. Exosome as a Novel Shuttle for Delivery of Therapeutics across Biological Barriers. Molecular Pharmaceutics. American Chemical Society January 7, 2019, pp 24– 40. 10.1021/acs.molpharmaceut.8b00901.

(31) Chen, C. C.; Liu, L.; Ma, F.; Wong, C. W.; Guo, X. E.; Chacko, J. V.; Farhoodi, H. P.; Zhang, S. X.; Zimak, J.; Ségaliny, A.; Riazifar, M.; Pham, V.; Digman, M. A.; Pone, E. J.; Zhao, W. Elucidation of Exosome Migration Across the Blood–Brain Barrier Model In Vitro. Cell. Mol. Bioeng. 2016, 9 (4), 509– 529. 10.1007/s12195-016-0458-3.

(32) Sáenz-Cuesta, M. Methods for Extracellular Vesicles Isolation in a Hospital Setting. Front. Immunol. 2015, 6 (FEB). 10.3389/fimmu.2015.00050.

(33) Rani, S.; O’Brien, K.; Kelleher, F. C.; Corcoran, C.; Germano, S.; Radomski, M. W.; Crown, J.; O’Driscoll, L. Isolation of Exosomes for Subsequent MRNA, MicroRNA, and Protein Profiling. In Methods in Molecular Biology; Humana Press, 2011; Vol. 784, pp 181–195. 10.1007/978-1-61779-289-2_13.

(34) Wiklander, O. P. B.; Nordin, J. Z.; O’Loughlin, A.; Gustafsson, Y.; Corso, G.; Mäger, I.; Vader, P.; Lee, Y.; Sork, H.; Seow, Y.; Heldring, N.; Alvarez-Erviti, L.; Smith, C. E.; Le Blanc, K.; Macchiarini, P.; Jungebluth, P.; Wood, M. J. A.; Andaloussi, S. EL. Extracellular Vesicle in Vivo Biodistribution Is Determined by Cell Source, Route of Administration and Targeting. J. Extracell. Vesicles 2015, 4 (1), 26316. 10.3402/jev.v4.26316.

(35) Debnath, K.; Las Heras, K.; Rivera, A.; Lenzini, S.; Shin, J.-W. Extracellular Vesicle–Matrix Interactions. Nat. Rev. Mater. 2023, 8 (6), 390–402. 10.1038/s41578-023-00551-3.

(36) Buzás, E. I.; Tóth, E. Á.; Sódar, B. W.; Szabó-Taylor, K. É. Molecular Interactions at the Surface of Extracellular Vesicles. Semin. Immunopathol. 2018, 40 (5), 453–464. 10.1007/s00281-018-0682-0.

(37) Palmulli, R.; Bresteau, E.; Raposo, G.; Montagnac, G.; van Niel, G. In Vitro Interaction of Melanoma-Derived Extracellular Vesicles with Collagen. Int. J. Mol. Sci. 2023, 24 (4), 3703. 10.3390/ijms24043703.

(38) Shimoda, M. Extracellular Vesicle-Associated MMPs: A Modulator of the Tissue Microenvironment. In Advances in Clinical Chemistry; Elsevier, 2019; Vol. 88, pp 35–66. 10.1016/bs.acc.2018.10.006.

(39) Arif, S.; Larochelle, S.; Trudel, B.; Gounou, C.; Bordeleau, F.; Brisson, A. R.; Moulin, V. J. The Diffusion of Normal Skin Wound Myofibroblast-derived Microvesicles Differs According to Matrix Composition. J. Extracell. Biol. 2024, 3 (1). 10.1002/jex2.131.

(40) Hoshino, A.; Costa-Silva, B.; Shen, T.-L.; Rodrigues, G.; Hashimoto, A.; Tesic Mark, M.; Molina, H.; Kohsaka, S.; Di Giannatale, A.; Ceder, S.; Singh, S.; Williams, C.; Soplop, N.; Uryu, K.; Pharmer, L.; King, T.; Bojmar, L.; Davies, A. E.; Ararso, Y.; Zhang, T.; Zhang, H.; Hernandez, J.; Weiss, J. M.; Dumont-Cole, V. D.; Kramer, K.; Wexler, L. H.; Narendran, A.; Schwartz, G. K.; Healey, J. H.; Sandstrom, P.; Jørgen Labori, K.; Kure, E. H.; Grandgenett, P. M.; Hollingsworth, M. A.; de Sousa, M.; Kaur, S.; Jain, M.; Mallya, K.; Batra, S. K.; Jarnagin, W. R.; Brady, M. S.; Fodstad, O.; Muller, V.; Pantel, K.; Minn, A. J.; Bissell, M. J.; Garcia, B. A.; Kang, Y.; Rajasekhar, V. K.; Ghajar, C. M.; Matei, I.; Peinado, H.; Bromberg, J.; Lyden, D. Tumour Exosome Integrins Determine Organotropic Metastasis. Nature 2015, 527 (7578), 329–335. 10.1038/nature15756.

(41) Hu, M.; Kenific, C. M.; Boudreau, N.; Lyden, D. Tumor-Derived Nanoseeds Condition the Soil for Metastatic Organotropism. Semin. Cancer Biol. 2023, 93, 70–82. 10.1016/j.semcancer.2023.05.003.

(42) Böing, A. N.; van der Pol, E.; Grootemaat, A. E.; Coumans, F. A. W.; Sturk, A.; Nieuwland, R. Single-step Isolation of Extracellular Vesicles by Size-exclusion Chromatography. J. Extracell. Vesicles 2014, 3 (1). 10.3402/jev.v3.23430.

(43) Benedikter, B. J.; Bouwman, F. G.; Vajen, T.; Heinzmann, A. C. A.; Grauls, G.; Mariman, E. C.; Wouters, E. F. M.; Savelkoul, P. H.; Lopez-Iglesias, C.; Koenen, R. R.; Rohde, G. G. U.; Stassen, F. R. M. Ultrafiltration Combined with Size Exclusion Chromatography Efficiently Isolates Extracellular Vesicles from Cell Culture Media for Compositional and Functional Studies. Sci. Rep. 2017, 7 (1), 15297. 10.1038/s41598-017-15717-7.

(44) Sbalzarini, I. F.; Koumoutsakos, P. Feature Point Tracking and Trajectory Analysis for Video Imaging in Cell Biology. J. Struct. Biol. 2005, 151 (2), 182–195. 10.1016/j.jsb.2005.06.002.

(45) González-King, H.; Tejedor, S.; Ciria, M.; Gil-Barrachina, M.; Soriano-Navarro, M.; Sánchez-Sánchez, R.; Sepúlveda, P.; García, N. A. Non-Classical Notch Signaling by MDA-MB-231 Breast Cancer Cell-Derived Small Extracellular Vesicles Promotes Malignancy in Poorly Invasive MCF-7 Cells. Cancer Gene Ther. 2022, 29 (7), 1056–1069. 10.1038/s41417-021-00411-8.

(46) Borgheti-Cardoso, L. N.; Kooijmans, S. A. A.; Chamorro, L. G.; Biosca, A.; Lantero, E.; Ramírez, M.; Avalos-Padilla, Y.; Crespo, I.; Fernández, I.; Fernandez-Becerra, C.; del Portillo, H. A.; Fernàndez-Busquets, X. Extracellular Vesicles Derived from Plasmodium-Infected and Non-Infected Red Blood Cells as Targeted Drug Delivery Vehicles. Int. J. Pharm. 2020, 587, 119627. 10.1016/j.ijpharm.2020.119627.

(47) Skotland, T.; Sandvig, K.; Llorente, A. Lipids in Exosomes: Current Knowledge and the Way Forward. Progress in Lipid Research. Elsevier Ltd April 1, 2017, pp 30–41. 10.1016/j.plipres.2017.03.001.

(48) Sezgin, E.; Kaiser, H.-J.; Baumgart, T.; Schwille, P.; Simons, K.; Levental, I. Elucidating Membrane Structure and Protein Behavior Using Giant Plasma Membrane Vesicles. Nat. Protoc. 2012, 7 (6), 1042–1051. 10.1038/nprot.2012.059.

(49) Steinkühler, J.; Fonda, P.; Bhatia, T.; Zhao, Z.; Leomil, F. S. C.; Lipowsky, R.; Dimova, R. Superelasticity of Plasma- and Synthetic Membranes Resulting from Coupling of Membrane Asymmetry, Curvature, and Lipid Sorting. Adv. Sci. 2021, 8 (21), 2102109. 10.1002/advs.202102109.

(50) Alter, C. L.; Detampel, P.; Schefer, R. B.; Lotter, C.; Hauswirth, P.; Puligilla, R. D.; Weibel, V. J.; Schenk, S. H.; Heusermann, W.; Schürz, M.; Meisner-Kober, N.; Palivan, C.; Einfalt, T.; Huwyler, J. High Efficiency Preparation of Monodisperse Plasma Membrane Derived Extracellular Vesicles for Therapeutic Applications. Commun. Biol. 2023, 6 (1), 478. 10.1038/s42003-023-04859-2.

(51) Iavello, A.; Frech, V. S. L.; Gai, C.; Deregibus, M. C.; Quesenberry, P. J.; Camussi, G. Role of Alix in MiRNA Packaging during Extracellular Vesicle Biogenesis. Int. J. Mol. Med. 2016, 37 (4), 958–966. 10.3892/ijmm.2016.2488.

(52) Romancino, D. P.; Buffa, V.; Caruso, S.; Ferrara, I.; Raccosta, S.; Notaro, A.; Campos, Y.; Noto, R.; Martorana, V.; Cupane, A.; Giallongo, A.; D’Azzo, A.; Manno, M.; Bongiovanni, A. Palmitoylation Is a Post-Translational Modification of Alix Regulating the Membrane Organization of Exosome-like Small Extracellular Vesicles. Biochim. Biophys. Acta - Gen. Subj. 2018, 1862 (12), 2879–2887. 10.1016/j.bbagen.2018.09.004.

(53) Chatellard-Causse, C.; Blot, B.; Cristina, N.; Torch, S.; Missotten, M.; Sadoul, R. Alix (ALG-2-Interacting Protein X), a Protein Involved in Apoptosis, Binds to Endophilins and Induces Cytoplasmic Vacuolization. J. Biol. Chem. 2002, 277 (32), 29108–29115. 10.1074/jbc.M204019200.

(54) Mathieu, M.; Névo, N.; Jouve, M.; Valenzuela, J. I.; Maurin, M.; Verweij, F. J.; Palmulli, R.; Lankar, D.; Dingli, F.; Loew, D.; Rubinstein, E.; Boncompain, G.; Perez, F.; Théry, C. Specificities of Exosome versus Small Ectosome Secretion Revealed by Live Intracellular Tracking of CD63 and CD9. Nat. Commun. 2021, 12 (1), 4389. 10.1038/s41467-021-24384-2.

(55) Logozzi, M.; De Milito, A.; Lugini, L.; Borghi, M.; Calabrò, L.; Spada, M.; Perdicchio, M.; Marino, M. L.; Federici, C.; Iessi, E.; Brambilla, D.; Venturi, G.; Lozupone, F.; Santinami, M.; Huber, V.; Maio, M.; Rivoltini, L.; Fais, S. High Levels of Exosomes Expressing CD63 and Caveolin-1 in Plasma of Melanoma Patients. PLoS One 2009, 4 (4), e5219. 10.1371/journal.pone.0005219.

(56) Kong, J. N.; He, Q.; Wang, G.; Dasgupta, S.; Dinkins, M. B.; Zhu, G.; Kim, A.; Spassieva, S.; Bieberich, E. Guggulsterone and Bexarotene Induce Secretion of Exosome-associated Breast Cancer Resistance Protein and Reduce Doxorubicin Resistance in <scp>MDA-MB</Scp> -231 Cells. Int. J. Cancer 2015, 137 (7), 1610–1620. 10.1002/ijc.29542.

(57) Metzelaar, M. J.; Wijngaard, P. L.; Peters, P. J.; Sixma, J. J.; Nieuwenhuis, H. K.; Clevers, H. C. CD63 Antigen. A Novel Lysosomal Membrane Glycoprotein, Cloned by a Screening Procedure for Intracellular Antigens in Eukaryotic Cells. J. Biol. Chem. 1991, 266 (5), 3239–3245. 10.1016/S0021-9258(18)49980-2.

(58) Sezgin, E.; Gutmann, T.; Buhl, T.; Dirkx, R.; Grzybek, M.; Coskun, Ü.; Solimena, M.; Simons, K.; Levental, I.; Schwille, P. Adaptive Lipid Packing and Bioactivity in Membrane Domains. PLoS One 2015, 10 (4), e0123930. 10.1371/journal.pone.0123930.

(59) Kaiser, H.-J.; Lingwood, D.; Levental, I.; Sampaio, J. L.; Kalvodova, L.; Rajendran, L.; Simons, K. Order of Lipid Phases in Model and Plasma Membranes. Proc. Natl. Acad. Sci. 2009, 106 (39), 16645–16650. 10.1073/pnas.0908987106.

(60) Gunther, G.; Malacrida, L.; Jameson, D. M.; Gratton, E.; Sánchez, S. A. LAURDAN since Weber: The Quest for Visualizing Membrane Heterogeneity. Acc. Chem. Res. 2021, 54 (4), 976– 987. 10.1021/acs.accounts.0c00687.

(61) Mangiarotti, A.; Siri, M.; Tam, N. W.; Zhao, Z.; Malacrida, L.; Dimova, R. Biomolecular Condensates Modulate Membrane Lipid Packing and Hydration. Nat. Commun. 2023, 14 (1), 6081. 10.1038/s41467-023-41709-5.

(62) Malacrida, L.; Astrada, S.; Briva, A.; Bollati-Fogolín, M.; Gratton, E.; Bagatolli, L. A. Spectral Phasor Analysis of LAURDAN Fluorescence in Live A549 Lung Cells to Study the Hydration and Time Evolution of Intracellular Lamellar Body-like Structures. Biochim. Biophys. Acta - Biomembr. 2016, 1858 (11), 2625–2635. 10.1016/j.bbamem.2016.07.017.

(63) Malacrida, L.; Ranjit, S.; Jameson, D. M.; Gratton, E. The Phasor Plot: A Universal Circle to Advance Fluorescence Lifetime Analysis and Interpretation. Annu. Rev. Biophys. 2021, 50 (1), 575–593. 10.1146/annurev-biophys-062920-063631.

(64) Aguilar, J.; Malacrida, L.; Gunther, G.; Torrado, B.; Torres, V.; Urbano, B. F.; Sánchez, S. A. Cells Immersed in Collagen Matrices Show a Decrease in Plasma Membrane Fluidity as the Matrix Stiffness Increases. Biochim. Biophys. Acta - Biomembr. 2023, 1865 (7), 184176. 10.1016/j.bbamem.2023.184176.

(65) Uppamoochikkal, P.; Tristram-Nagle, S.; Nagle, J. F. Orientation of Tie-Lines in the Phase Diagram of DOPC/DPPC/Cholesterol Model Biomembranes. Langmuir 2010, 26 (22), 17363–17368. 10.1021/la103024f.

(66) Steinkühler, J.; Sezgin, E.; Urbančič, I.; Eggeling, C.; Dimova, R. Mechanical Properties of Plasma Membrane Vesicles Correlate with Lipid Order, Viscosity and Cell Density. Commun. Biol. 2019, 2 (1), 337. 10.1038/s42003-019-0583-3.

(67) Sinn, C. G.; Antonietti, M.; Dimova, R. Binding of Calcium to Phosphatidylcholine– Phosphatidylserine Membranes. Colloids Surfaces A Physicochem. Eng. Asp. 2006, 282–283, 410–419. 10.1016/j.colsurfa.2005.10.014.

(68) Wess, T. J. Collagen Fibril Form and Function. In Advances in Protein Chemistry; Academic Press, 2005; Vol. 70, pp 341–374. 10.1016/S0065-3233(05)70010-3.

(69) Fratzl, P. Collagen: Structure and Mechanics, an Introduction. In Collagen; Springer US: Boston, MA, 2008; pp 1–13. 10.1007/978-0-387-73906-9_1.

(70) Jokinen, J.; Dadu, E.; Nykvist, P.; Käpylä, J.; White, D. J.; Ivaska, J.; Vehviläinen, P.; Reunanen, H.; Larjava, H.; Häkkinen, L.; Heino, J. Integrin-Mediated Cell Adhesion to Type I Collagen Fibrils. J. Biol. Chem. 2004, 279 (30), 31956–31963. 10.1074/jbc.M401409200.

(71) Gasperini, L.; Mano, J. F.; Reis, R. L. Natural Polymers for the Microencapsulation of Cells. J. R. Soc. Interface 2014, 11 (100), 20140817. 10.1098/rsif.2014.0817.

(72) Kaufman, L. J.; Brangwynne, C. P.; Kasza, K. E.; Filippidi, E.; Gordon, V. D.; Deisboeck, T. S.; Weitz, D. A. Glioma Expansion in Collagen I Matrices: Analyzing Collagen Concentration-Dependent Growth and Motility Patterns. Biophys. J. 2005, 89 (1), 635–650. 10.1529/biophysj.105.061994.

(73) Yang, Y.; Kaufman, L. J. Rheology and Confocal Reflectance Microscopy as Probes of Mechanical Properties and Structure during Collagen and Collagen/Hyaluronan Self-Assembly. Biophys. J. 2009, 96 (4), 1566–1585. 10.1016/j.bpj.2008.10.063.

(74) Tam, N. W.; Schullian, O.; Cipitria, A.; Dimova, R. Nonspecific Membrane-Matrix Interactions Influence Diffusivity of Lipid Vesicles in Hydrogels. Biophys. J. 2024, 123 (5), 638–650. 10.1016/j.bpj.2024.02.005.

(75) Lieleg, O.; Baumgärtel, R. M.; Bausch, A. R. Selective Filtering of Particles by the Extracellular Matrix: An Electrostatic Bandpass. Biophys. J. 2009, 97 (6), 1569–1577. 10.1016/j.bpj.2009.07.009.

(76) Xu, Q.; Ensign, L. M.; Boylan, N. J.; Schön, A.; Gong, X.; Yang, J.-C.; Lamb, N. W.; Cai, S.; Yu, T.; Freire, E.; Hanes, J. Impact of Surface Polyethylene Glycol (PEG) Density on Biodegradable Nanoparticle Transport in Mucus Ex Vivo and Distribution in Vivo. ACS Nano 2015, 9 (9), 9217–9227. 10.1021/acsnano.5b03876.

(77) Smyth, D. G.; Blumenfeld, O. O.; Konigsberg, W. Reactions of N-Ethylmaleimide with Peptides and Amino Acids. Biochem. J. 1964, 91 (3), 589–595. 10.1042/bj0910589.

(78) Levental, I.; Lingwood, D.; Grzybek, M.; Coskun, Ü.; Simons, K. Palmitoylation Regulates Raft Affinity for the Majority of Integral Raft Proteins. Proc. Natl. Acad. Sci. 2010, 107 (51), 22050–22054. 10.1073/pnas.1016184107.

(79) Del Piccolo, N.; Placone, J.; He, L.; Agudelo, S. C.; Hristova, K. Production of Plasma Membrane Vesicles with Chloride Salts and Their Utility as a Cell Membrane Mimetic for Biophysical Characterization of Membrane Protein Interactions. Anal. Chem. 2012, 84 (20), 8650–8655. 10.1021/ac301776j.

(80) Sakai-Kato, K.; Yoshida, K.; Takechi-Haraya, Y.; Izutsu, K. Physicochemical Characterization of Liposomes That Mimic the Lipid Composition of Exosomes for Effective Intracellular Trafficking. Langmuir 2020, 36 (42), 12735–12744. 10.1021/acs.langmuir.0c02491.

(81) Lemmon, M. A. Membrane Recognition by Phospholipid-Binding Domains. Nat. Rev. Mol. Cell Biol. 2008, 9 (2), 99–111. 10.1038/nrm2328.

(82) Melcrová, A.; Pokorna, S.; Pullanchery, S.; Kohagen, M.; Jurkiewicz, P.; Hof, M.; Jungwirth, P.; Cremer, P. S.; Cwiklik, L. The Complex Nature of Calcium Cation Interactions with Phospholipid Bilayers. Sci. Rep. 2016, 6 (1), 38035. 10.1038/srep38035.

(83) Marsh, D.; Bartucci, R.; Sportelli, L. Lipid Membranes with Grafted Polymers: Physicochemical Aspects. Biochim. Biophys. Acta - Biomembr. 2003, 1615 (1–2), 33–59. 10.1016/S0005-2736(03)00197-4.

(84) Lee, H.; Larson, R. G. Adsorption of Plasma Proteins onto PEGylated Lipid Bilayers: The Effect of PEG Size and Grafting Density. Biomacromolecules 2016, 17 (5), 1757–1765. 10.1021/acs.biomac.6b00146.

(85) Du, H.; Chandaroy, P.; Hui, S. W. Grafted Poly-(Ethylene Glycol) on Lipid Surfaces Inhibits Protein Adsorption and Cell Adhesion. Biochim. Biophys. Acta - Biomembr. 1997, 1326 (2), 236–248. 10.1016/S0005-2736(97)00027-8.

(86) Maneewongvatana, S.; Mount, D. M. It’s Okay to Be Skinny, If Your Friends Are Fat. Cent. Geom. Comput. 4th Annu. Work. Comput. Geom. 1999, 2 (October), 1–8.

(87) Buerkle, M. A.; Pahernik, S. A.; Sutter, A.; Jonczyk, A.; Messmer, K.; Dellian, M. Inhibition of the Alpha-? Integrins with a Cyclic RGD Peptide Impairs Angiogenesis, Growth and Metastasis of Solid Tumours in Vivo. Br. J. Cancer 2002, 86 (5), 788–795. 10.1038/sj.bjc.6600141.

(88) Meena, C. L.; Singh, D.; Weinmüller, M.; Reichart, F.; Dangi, A.; Marelli, U. K.; Zahler, S.; Sanjayan, G. J. Novel Cilengitide-Based Cyclic RGD Peptides as ?vβ Integrin Inhibitors. Bioorg. Med. Chem. Lett. 2020, 30 (8), 127039. 10.1016/j.bmcl.2020.127039.

(89) Russo, M. A.; Paolillo, M.; Sanchez-Hernandez, Y.; Curti, D.; Ciusani, E.; Serra, M.; Colombo, L.; Schinelli, S. A Small-Molecule RGD-Integrin Antagonist Inhibits Cell Adhesion, Cell Migration and Induces Anoikis in Glioblastoma Cells. Int. J. Oncol. 2013, 42 (1), 83–92. 10.3892/ijo.2012.1708.

(90) Kapp, T. G.; Rechenmacher, F.; Neubauer, S.; Maltsev, O. V.; Cavalcanti-Adam, E. A.; Zarka, R.; Reuning, U.; Notni, J.; Wester, H.-J.; Mas-Moruno, C.; Spatz, J.; Geiger, B.; Kessler, H. A Comprehensive Evaluation of the Activity and Selectivity Profile of Ligands for RGD-Binding Integrins. Sci. Rep. 2017, 7 (1), 39805. 10.1038/srep39805.

(91) Takahashi, S.; Leiss, M.; Moser, M.; Ohashi, T.; Kitao, T.; Heckmann, D.; Pfeifer, A.; Kessler, H.; Takagi, J.; Erickson, H. P.; Fässler, R. The RGD Motif in Fibronectin Is Essential for Development but Dispensable for Fibril Assembly. J. Cell Biol. 2007, 178 (1), 167–178. 10.1083/jcb.200703021.

(92) Barczyk, M.; Carracedo, S.; Gullberg, D. Integrins. Cell Tissue Res. 2010, 339 (1), 269–280. 10.1007/s00441-009-0834-6.

(93) Davis, G. E. Affinity of Integrins for Damaged Extracellular Matrix: ?vβ3 Binds to Denatured Collagen Type I through RGD Sites. Biochem. Biophys. Res. Commun. 1992, 182 (3), 1025–1031. 10.1016/0006-291X(92)91834-D.

(94) Taubenberger, A. V.; Woodruff, M. A.; Bai, H.; Muller, D. J.; Hutmacher, D. W. The Effect of Unlocking RGD-Motifs in Collagen I on Pre-Osteoblast Adhesion and Differentiation. Biomaterials 2010, 31 (10), 2827–2835. 10.1016/j.biomaterials.2009.12.051.

(95) Ramanujan, S.; Pluen, A.; McKee, T. D.; Brown, E. B.; Boucher, Y.; Jain, R. K. Diffusion and Convection in Collagen Gels: Implications for Transport in the Tumor Interstitium. Biophys. J. 2002, 83 (3), 1650–1660. 10.1016/S0006-3495(02)73933-7.

(96) Emsley, J.; Knight, C. G.; Farndale, R. W.; Barnes, M. J.; Liddington, R. C. Structural Basis of Collagen Recognition by Integrin ?2β1. Cell 2000, 101 (1), 47–56. 10.1016/S0092-8674(00)80622-4.

(97) Knight, C. G.; Morton, L. F.; Peachey, A. R.; Tuckwell, D. S.; Farndale, R. W.; Barnes, M. J. The Collagen-Binding A-Domains of Integrins ?1β1 and ?2β1Recognize the Same Specific Amino Acid Sequence, GFOGER, in Native (Triple-Helical) Collagens. J. Biol. Chem. 2000, 275 (1), 35–40. 10.1074/jbc.275.1.35.

(98) Evnin, L. B.; Vásquez, J. R.; Craik, C. S. Substrate Specificity of Trypsin Investigated by Using a Genetic Selection. Proc. Natl. Acad. Sci. 1990, 87 (17), 6659–6663. 10.1073/pnas.87.17.6659.

(99) Kim, C.-S.; Yoo, H. Three-Dimensional Confocal Reflectance Microscopy for Surface Metrology. Meas. Sci. Technol. 2021, 32 (10), 102002. 10.1088/1361-6501/ac04df.

(100) Chen, X.; Nadiarynkh, O.; Plotnikov, S.; Campagnola, P. J. Second Harmonic Generation Microscopy for Quantitative Analysis of Collagen Fibrillar Structure. Nat. Protoc. 2012, 7 (4), 654– 669. 10.1038/nprot.2012.009.

(101) Brightman, A. O.; Rajwa, B. P.; Sturgis, J. E.; McCallister, M. E.; Robinson, J. P.; Voytik-Harbin, S. L. Time-Lapse Confocal Reflection Microscopy of Collagen Fibrillogenesis and Extracellular Matrix Assembly in Vitro. Biopolymers 2000, 54 (3), 222– 234. 10.1002/1097-0282(200009)54:3<222::AID-BIP80>3.0.CO;2-K.

(102) Condeelis, J.; Segall, J. E. Intravital Imaging of Cell Movement in Tumours. Nat. Rev. Cancer 2003, 3 (12), 921–930. 10.1038/nrc1231.

(103) Hyenne, V.; Ghoroghi, S.; Collot, M.; Bons, J.; Follain, G.; Harlepp, S.; Mary, B.; Bauer, J.; Mercier, L.; Busnelli, I.; Lefebvre, O.; Fekonja, N.; Garcia-Leon, M. J.; Machado, P.; Delalande, F.; López, A. A.; Silva, S. G.; Verweij, F. J.; van Niel, G.; Djouad, F.; Peinado, H.; Carapito, C.; Klymchenko, A. S.; Goetz, J. G. Studying the Fate of Tumor Extracellular Vesicles at High Spatiotemporal Resolution Using the Zebrafish Embryo. Dev. Cell 2019, 48 (4), 554-572.e7. 10.1016/j.devcel.2019.01.014.

(104) Atchison, D. K.; Beierwaltes, W. H. The Influence of Extracellular and Intracellular Calcium on the Secretion of Renin. Pflügers Arch. - Eur. J. Physiol. 2013, 465 (1), 59–69. 10.1007/s00424-012-1107-x.

