## Supplemental Materials for "Extracellular vesicle mobility in collagen I hydrogels is influenced by RGD-binding integrins"

Nicky W. Tam[1]*, Alexander Becker[1,2], Agustín Mangiarotti[1], Amaia Cipitria [1,3,4]*, and Rumiana Dimova[1]*

[1]Max Planck Institute of Colloids and Interfaces, Science Park Golm, 14476 Potsdam, Germany

[2]McGill University, Montréal, Canada, H3A 0G4

[3]Group of Bioengineering in Regeneration and Cancer, Biogipuzkoa Health Research Institute, San Sebastián, 20014 Spain

[4]IKERBASQUE, Basque Foundation for Science, 48009 Bilbao, Spain

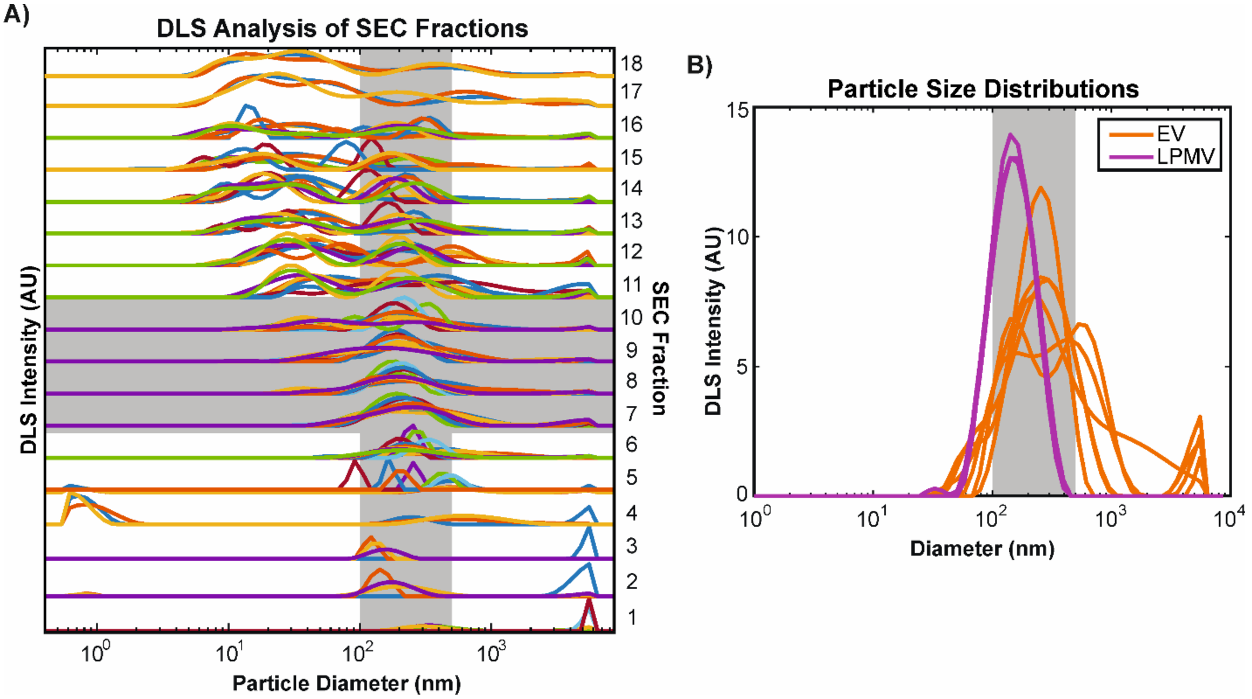


Supplemental Figure S1, DLS analysis of EV and LPMV size distribution. A) Size distributions of particles detected by DLS in the different fractions collected during SEC purification of EVs. Different colours represent different replicates. The target particle size range of 100-400nm and the collected fractions are shaded in grey to show that they intersect. Y-axes of the size distributions represent DLS intensity and have been normalized to show relative enrichment as opposed to absolute abundance. Although fraction 6 appears to have EVs in the target size range, the overall particle enrichment is much lower than in fractions 7-10. B) Comparison of EV (orange) and LPMV (magenta) size distributions. EV traces represent pooled EV fractions. EVs are expected to be more polydisperse because LPMVs are extruded with a defined filter pore size. Different traces represent different replicates. The shaded grey area corresponds to the target size range of 100-400nm.

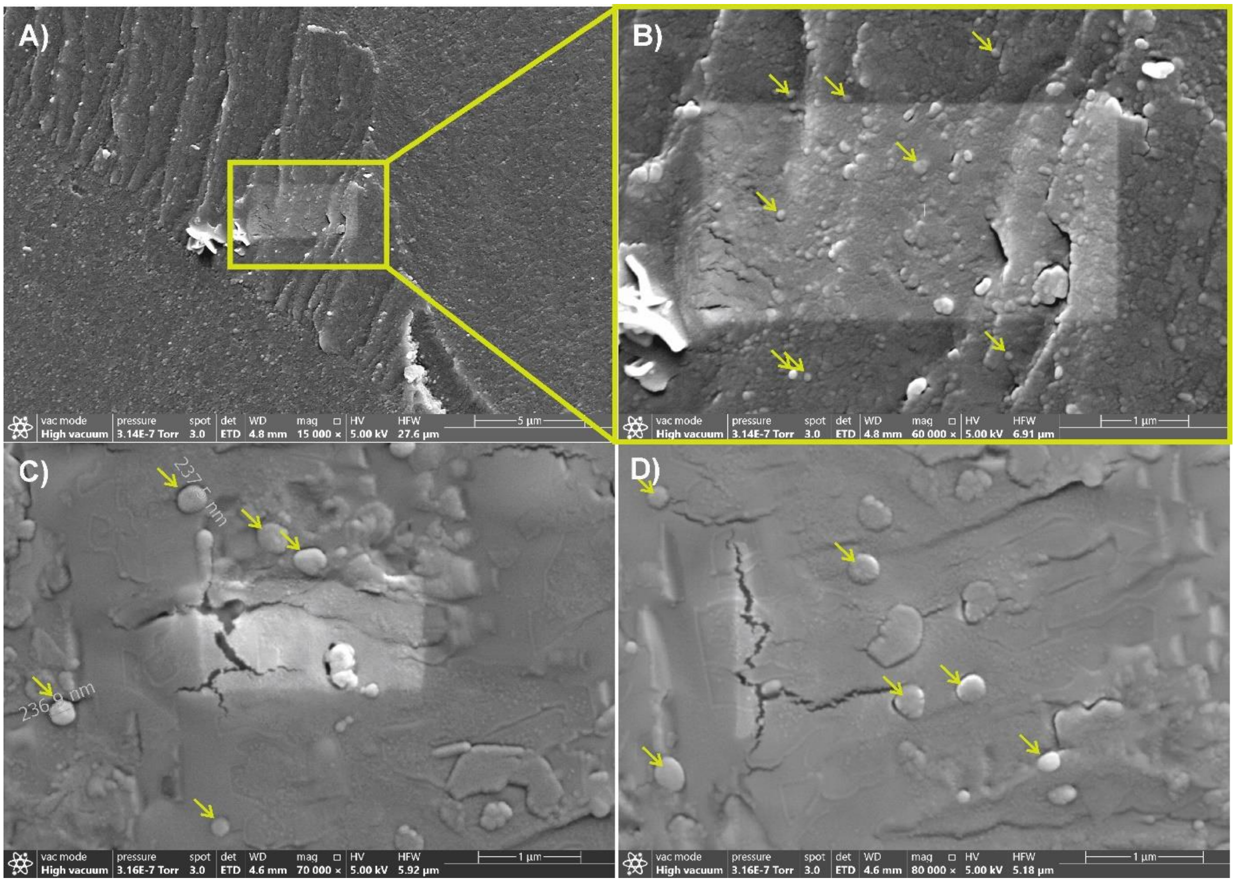


Supplemental Figure S2. A) Raw CryoSEM image of EVs captured at 15000× magnification. EVs have been compacted by concentrating the sample with a centrifugal filter and appear as a grainy mass on the freeze-fractured surface. B) Raw CryoSEM image of EVs captured at 60000× magnification. Image corresponds the central region of A, as indicated. Examples of EVs are indicated with yellow arrows. C) CryoSEM image of EVs at 70000× magnification. In this sample, EVs appear more dispersed than in Figures S2 and S3, partially or fully embedded in the surrounding frozen medium. Overlays show the measured diameters of 2 identified EVs. D) CryoSEM image of EVs at 80000× magnification. Here, some of the EVs appear to have been cleaved in half during freeze fracture, revealing a flat, cross-sectional surface.

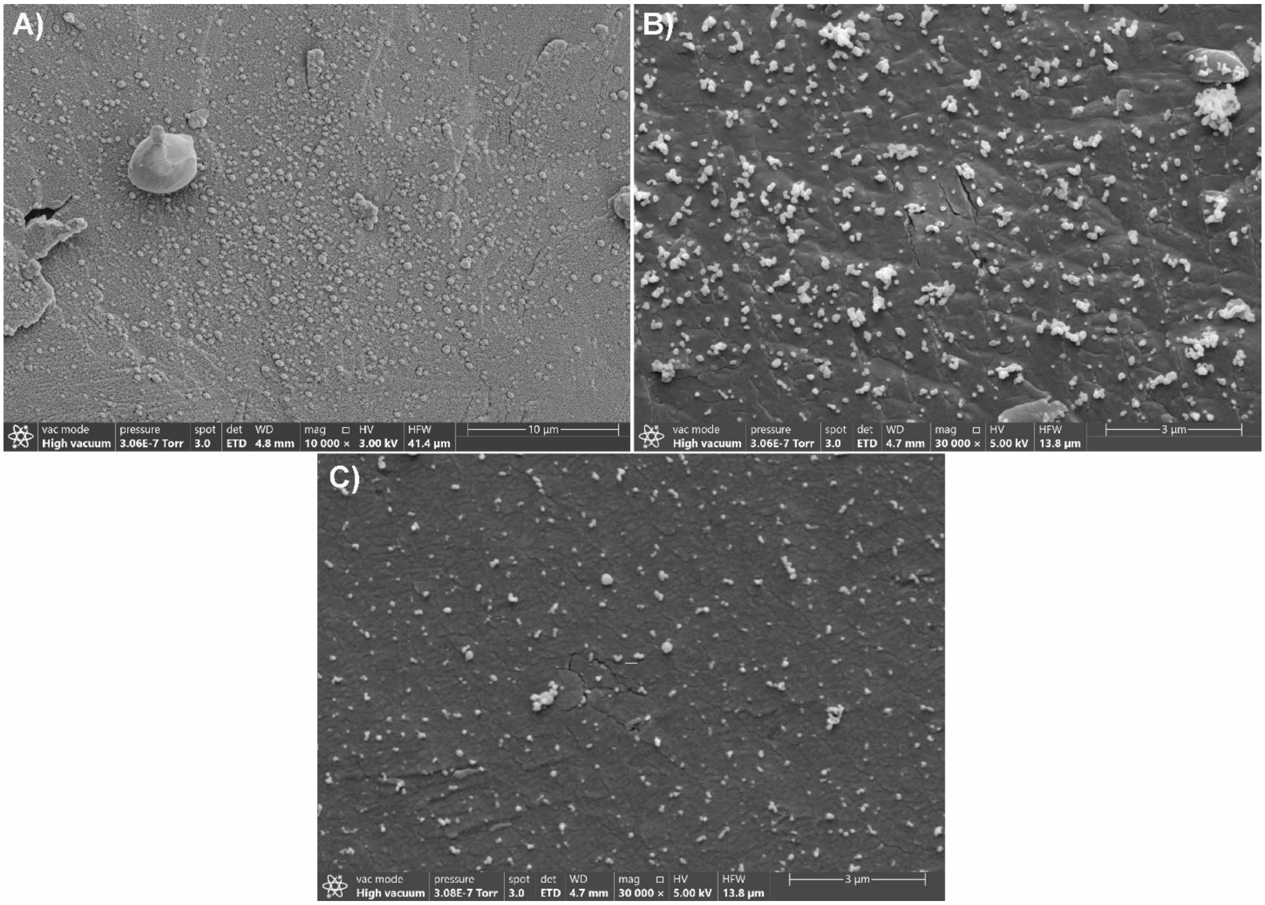


Supplemental Figure S3, A) Raw CryoSEM image of LPMVs at 10000× magnification. LPMVs were also compacted during concentration with centrifugal filters, but not to the same extent as the EVs in Figure S2. B) Raw CryoSEM image of LPMVs at 30000× magnification. Here, LPMVs are found in small aggregates of varying size, likely formed during concentration with centrifugal filters. C) Raw CryoSEM image of LPMVs at 30000× magnification. In this sample, LPMVs are well-dispersed with only a few aggregates.

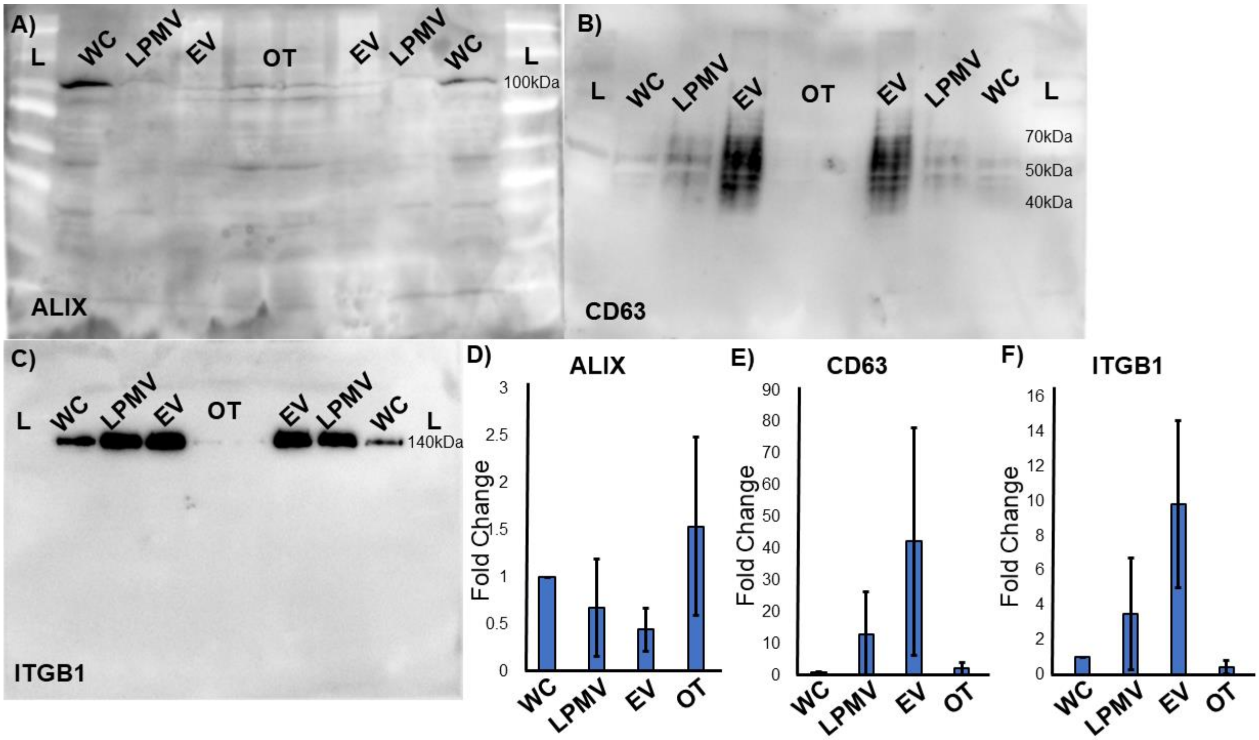


Supplemental Figure S4, Western blot analysis of protein expression. A-C) The full uncropped blots from Fig. 2B in the main text are presented here for reference. Samples were loaded, from left to right: ladder (L), whole cell lysate (WC), LPMVs, EVs, non-EV off-target SEC fractions (OT), non EV off-target SEC fractions, EVs, LPMVs, whole cell lysate, ladder. The same blot was stripped and re-probed twice to get a direct comparison of protein enrichment. A) ALIX expression appears at the expected size of 100kDa with some off-target binding. B) CD63 appears with broad smearing between 40 and 70kDa with little apparent off-target binding. C) integrin β-1 is clearly expressed as a single, well-defined band at 140kDa with no off-target or non-specific binding. D-F) Densitometric analysis of Western blot data from n=6 replicates. Data show integrated pixel density of blots, as determined with ImageJ, normalized to the whole cell lysate in each blot as a control. ALIX expression appears to be inversely proportional to CD63 and ITGB1 expression.

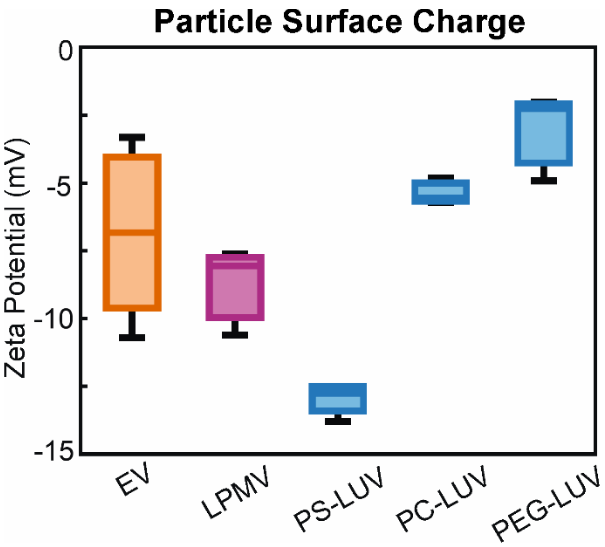


Supplemental Figure S5, Comparison of particle surface charges, measured as zeta potential. EVs are compared with LPMVs and LUVs composed of pure DOPC (PC-LUV), 4:1 DOPC+DOPS (PS-LUV), and DOPC+1mol% DSPE-mPEG1K (PEG-LUV). Zeta potential values were measured in calcium-free high salt-concentration buffer (HBS) and thus may be affected by charge screening. Nevertheless, values can be taken as relative measures of surface charge. It should be noted that although PS-LUVs were produced with 20% phosphatidylserine to reflect the composition of EVs, as previously found in lipidomics studies, the discrepancy in surface charge between EVs and PS-LUVs should be expected. EV membranes are crowded with proteins and intact glycocalyx, which can alter overall surface charge. Sample sizes are n=7 for EVs and n=3 for all other particles.

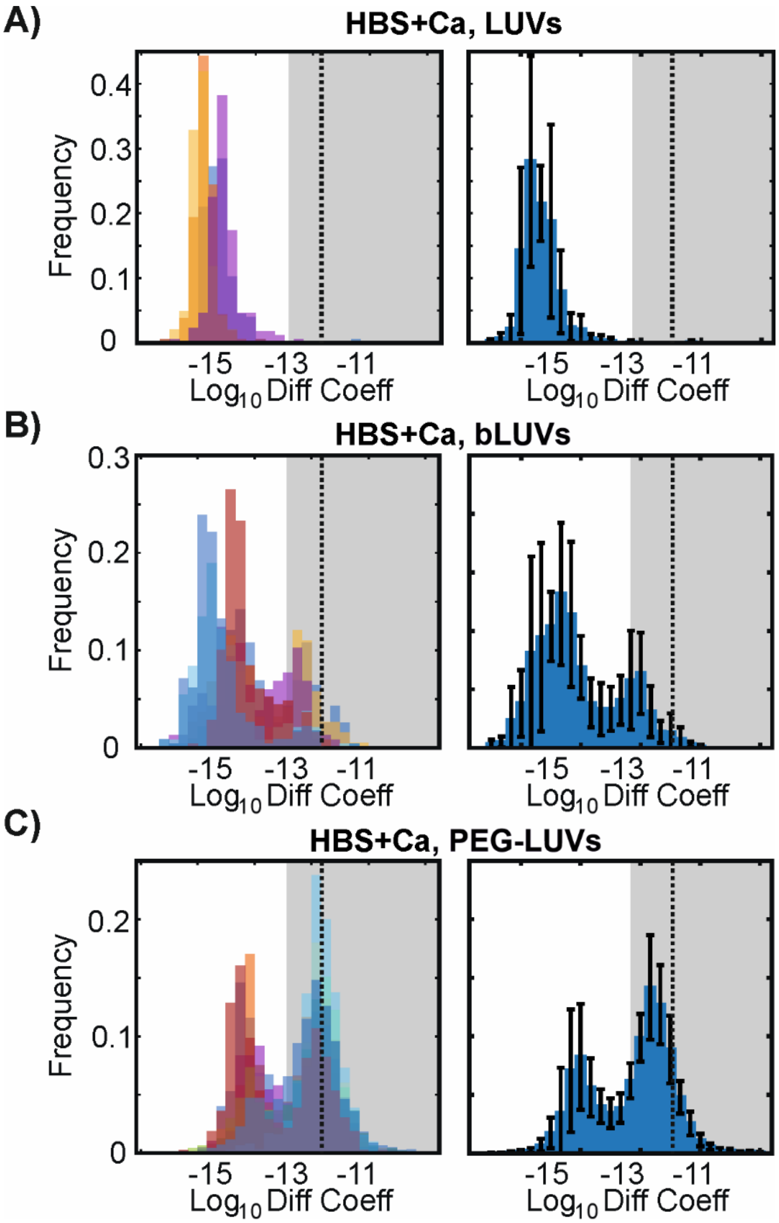


Supplemental Figure S6, Histograms of $\log_{10}$ diffusion coefficients. Individual experimental replicates are shown in the left column in different colours and the mean distributions are shown in the right column with error bars showing standard deviation. Grey shaded areas show the mobile fraction above the mobility threshold of -13. A dotted line shows the Stokes-Einstein predicted value for an ideal 200nm-diameter spherical particle diffusing in liquid water (-12). A) DOPC LUVs are entirely immobilized in the collagen I hydrogels. B) DOPC LUVs previously incubated with solubilized collagen I to 'block' their surfaces (bLUVs) regain some mobility in hydrogels. C) PEGylated LUVs (PEG-LUVs) show the greatest mobility of the synthetic membranes tested, and have a similar mobile fraction to EVs. They show, however, more polarized behaviour, with particles either fully immobilized or very mobile, with fewer particles with intermediate diffusion coefficient values. All LUVs were tested in calcium-containing buffer (HBS+Ca).

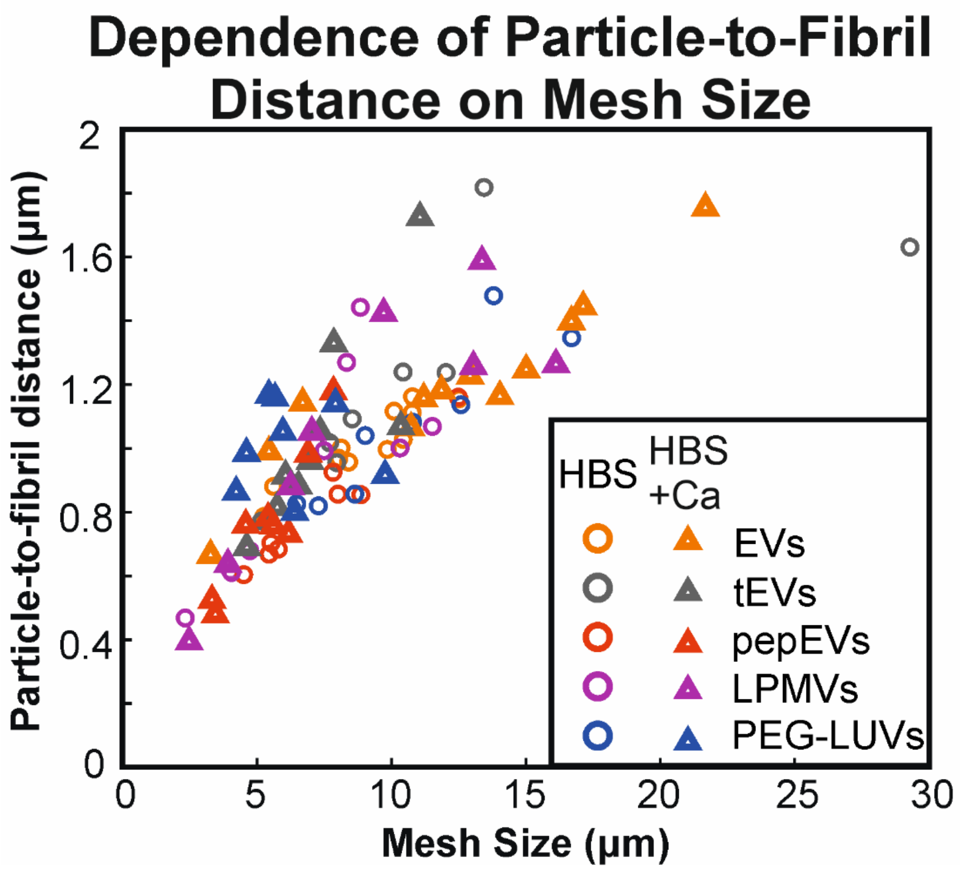


Supplemental Figure S7, Relationship between hydrogel mesh size and particle-to-fibril distance. Each data point represents a different replicate, with the colours showing the different particles tested in calcium-containing (HBS+Ca, triangles) and calcium-free (HBS, circles) buffer. Mesh size was not varied on purpose, but arose by natural heterogeneity between hydrogel samples. Average particle-to-fibril distances of each sample were normalized by the sample mesh size, which was determined in parallel. This removed the mesh size dependence and decreased overall variance in the data set. Alternatively, the normalized data can also be interpreted as the slope of the trend between particle-to-fibril distance and mesh size.

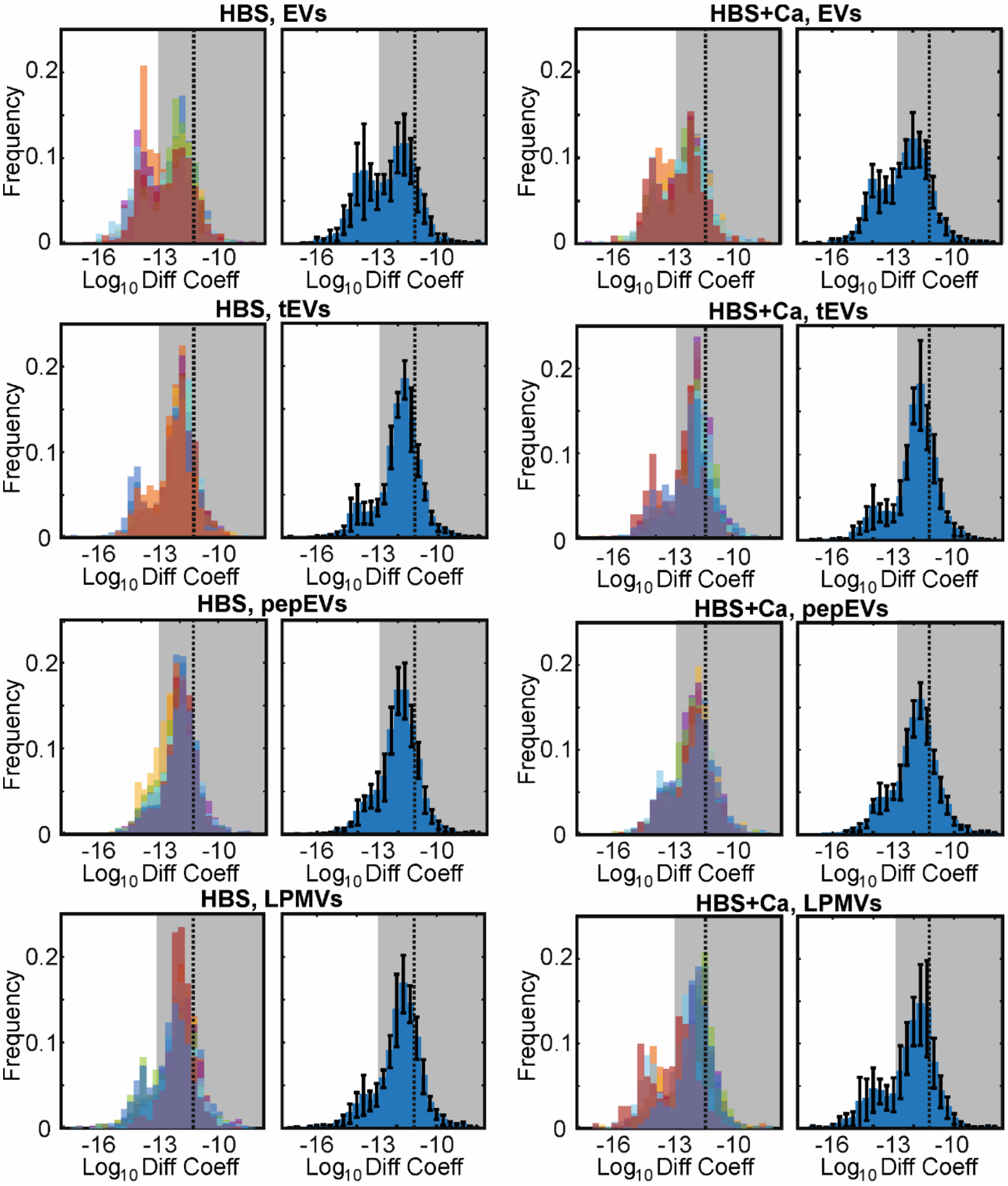


S

Supplemental Figure S8, Distributions of $log_{10}$ diffusion coefficients of cell-derived vesicles diffusing in collagen I hydrogels with calcium-free (HBS) and calcium-containing (HBS+Ca) buffers. Histograms with different colours show individual experimental replicates (n=6 each), while blue histograms with error bars show mean distributions with standard deviation. A dotted line indicates the threshold value of -13 used to determine mobile fraction.

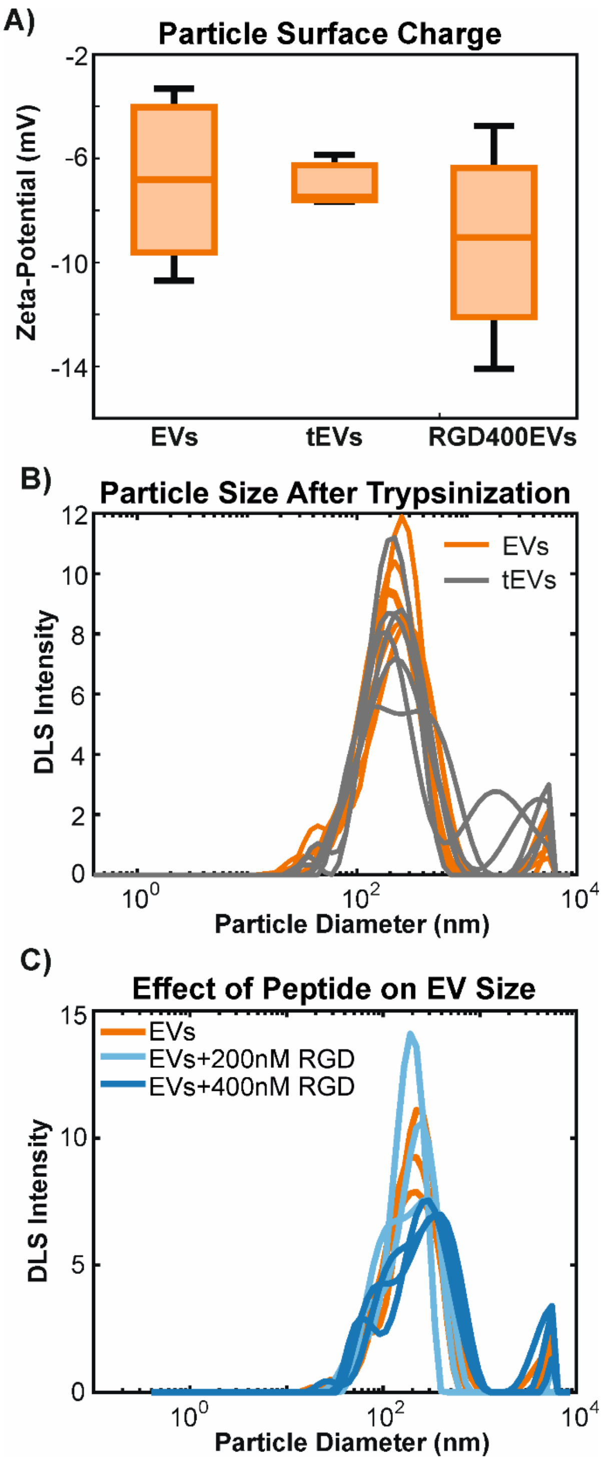


Supplemental Figure S9, Effect of trypsinization and RGD peptide on EV surface charge and size. A) Relative surface charges of EVs under different treatments, as represented by zeta-potential measurements in high ionic strength buffers (HBS). Trypsinization (tEVs) and treatment with 400 nM RGD peptide (RGD400EVs) do not appear to affect EV surface charge. Sample sizes are n=7 for control EVs and n=3 for other particles. B) Size distributions of EVs before (orange) and after (grey) trypsinization. Individual traces represent separate experimental replicates (n=6 for each condition). While overall size does not appear to change much, there appears to be slightly higher variability in size and the possible existence of aggregates in tEVs (seen here as extra peaks). C) Effect of RGD peptide on EV size distribution. EV size distribution does not appear to be greatly affected by treatment with 200 nM peptide (light blue), but the distribution begins to appear broader and multiple peaks can be seen at 400 nM peptide (dark blue), suggesting the formation of EV aggregates. Individual traces represent different replicates with sample sizes of n=3 for each condition.

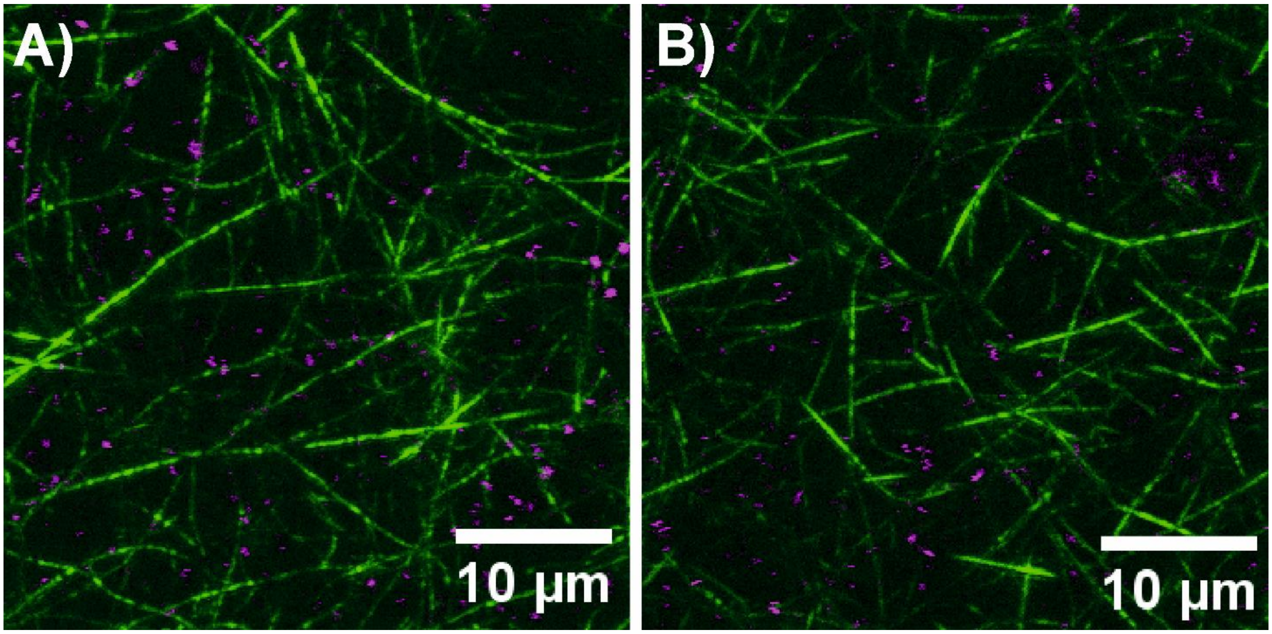


Supplemental Figure S10, Representative examples of EVs diffusing in collagen I hydrogels. Images consist of compound vertical projections of confocal reflectance images of collagen I fibrils (green) and confocal fluorescence microscopy of fluorescently-labelled EVs in calcium-free HBS (A) and calcium-containing HBS+Ca (B). Vertical projection was done over a stack of 10 images with 0.75µm spacing between slices in order to capture more fibrils and particles than could be done with a single slice. Image analysis for mesh size, particle-to-fibril distance, and fibril association were nonetheless conducted on single slices and averaged over the whole stack.
